# Intracellular diffusion in the cytoplasm increases with cell size in fission yeast

**DOI:** 10.1101/2024.09.21.613766

**Authors:** Catherine Tan, Michael C. Lanz, Matthew Swaffer, Jan Skotheim, Fred Chang

## Abstract

Diffusion in the cytoplasm can greatly impact cellular processes, yet regulation of macromolecular diffusion remains poorly understood. There is increasing evidence that cell size affects the density and macromolecular composition of the cytoplasm. Here, we studied whether cell size affects diffusion at the scale of macromolecules tens of microns in diameter. We analyzed the diffusive motions of intracellular genetically-encoded multimeric 40 nm nanoparticles (cytGEMs) in the cytoplasm of the fission yeast *Schizosaccharomyces pombe*. Using cell size mutants, we showed that cytGEMs diffusion coefficients decreased in smaller cells and increased in larger cells. This increase in diffusion in large cells may be due to a decrease in the DNA-to-Cytoplasm ratio, as diffusion was not affected in large multinucleate cytokinesis mutants. In investigating the underlying causes of altered cytGEMs diffusion, we found that the proteomes of large and small cells exhibited size-specific changes, including the sub-scaling of ribosomal proteins in large cells. Comparison with a similar dataset from human cells revealed that features of size-dependent proteome remodeling were conserved. These studies demonstrate that cell size is an important parameter in determining the biophysical properties and the composition of the cytoplasm.

## INTRODUCTION

Cell size is an intrinsic physical property of all cells that can impact physiology from the cellular level to the organismal. Although cell size can vary over six orders of magnitude among diverse cell types, cell size varies within a much narrower range for a specific cell type due to homeostatic mechanisms (Ginzberg *et al*., 2015; Zatulovskiy and Skotheim, 2020). Cell size can impart different cellular functions and developmental potential (Hecht *et al*., 2016; Lengefeld *et al*., 2021). Aberrant cell size can also signify biological dysfunction and has been associated with aging, senescence, numerous cancers, and other human diseases (Lloyd, 2013). The mechanisms for how cell size impacts cellular physiology, however, remain poorly understood.

Recent studies have begun to implicate effects of cell size on the global properties of the cytoplasm. The cytoplasm can be regarded as a heterogenous, dynamic, and crowded viscoelastic matrix that exerts osmotic forces, and impacts nearly all biochemical reactions through effects on viscosity, macromolecular crowding, phase separation, and likely many other biophysical phenomena (Zhou *et al*., 2008; Mitchison, 2019). For instance, variations in density, which can be the result of complex dynamics between biosynthesis, degradation, and osmotic water fluxes, can cause significant changes in cellular physiology and/or function as seen during the cell cycle, differentiation, and stress (Neurohr and Amon, 2020).

Generally, the concentrations of cellular components are thought to be maintained at different cell sizes by scaling relationships. For instance, mRNA, protein, transcription, translation, and the volume of many organelles can scale with cell size (Elliott *et al*., 1979; Creanor and Mitchison, 1982; Elliott, 1983; Neumann and Nurse, 2007; Zhurinsky *et al*., 2010; Padovan-Merhar *et al*., 2015; Chadwick *et al*., 2020; Marshall, 2020; Basier and Nurse, 2023; Swaffer *et al*., 2023). However, such scaling relationships have limitations, and so a breakdown in scaling mechanisms could explain cell size-dependent changes in cellular physiology. For example, it was observed in budding yeast cells that were arrested in G1 phase and grew to very large sizes (up to 10 times larger in volume than normal) exhibited defects in protein synthesis, and progressively became less dense (Neurohr *et al*., 2019). Similar dilution effects have been seen in senescent metazoan cells, indicating that a large cell size may be causal for aspects of senescent physiology, and is not merely a side effect of senescence (Demidenko and Blagosklonny, 2008; Neurohr *et al*., 2019; Lengefeld *et al*., 2021; Lanz *et al*., 2022). Similarly, in fission yeast, cells arrested in G2 phase grow very large and show a gradual slowdown in rates of cell growth and protein translation, further illustrating that a large cell size is detrimental for proliferation and normal cell function (Knapp *et al*., 2019; Basier and Nurse, 2023). Recent studies have demonstrated that cell size can also change the composition of the proteome even in cases where overall protein concentration remains relatively constant between sizes (Schmoller et al., 2015; Keifenheim et al., 2017; Lans et al., 2022, 2024).

One proposed explanation for certain cellular pathologies associated with overly large cells is that the DNA-to-Cytoplasm ratio in these cells has dropped below a critical threshold required to scale biosynthetic processes (Zhurinsky *et al*., 2010; Marguerat and Bähler, 2012; Neurohr *et al*., 2019; Balachandra *et al*., 2022; Cadart and Heald, 2022; Xie *et al*., 2022). As cells grows larger without a concomitant increase in DNA, there may be insufficient gene copies or transcriptional or translational machinery to support biomass production for an exponentially-growing cell volume. This theory could explain why cells with increased ploidy can grow to larger sizes without exhibiting defects associated with large cell size (Neurohr *et al*., 2019; Mu *et al*., 2020; Lanz *et al*., 2022, 2024).

The fission yeast *S. pombe* is a leading model organism in defining cell size control mechanisms and scaling relationships (Nurse, 1985; Wood and Nurse, 2015). Many cell-size scaling studies utilize well characterized cell size mutants as such as *wee1-50* and *cdc25-22* mutants, which alter cell size by affecting the length of G2 phase through regulation of the CDK1 cell cycle dependent kinase (Nurse, 1975; Fantes and Nurse, 1978; Neumann and Nurse, 2007; Zhurinsky *et al*., 2010; Knapp *et al*., 2019; Pickering *et al*., 2019; Sun *et al*., 2020). Thus, the molecular bases for the perturbations in cell size and genomic copy number are generally well-defined. Another feature of fission yeast is that although some effects of cell size in some cell types may be caused changes in surface area-to-volume (SA/V) ratio (Harris *et al*., 2018), SA/V ratio varies little with size in fission yeast due to their characteristic rod-cell shape (Shi *et al*., 2021). Recent studies in fission yeast demonstrate how the intracellular density of the cytoplasm fluctuates during the cell cycle and how properties of the cytoplasm are altered in starvation responses and during sporulation (Joyner *et al*., 2016; Munder *et al*., 2016; Heimlicher *et al*., 2019; Odermatt *et al*., 2021; Sakai *et al*., 2024). However, it remains unclear which cellular components are responsible for fluctuations in cytoplasmic density or how these fluctuations alter cellular functions when cell size changes.

Here, we studied the effects of cell size on the biophysical properties of the fission yeast cytoplasm by assessing the diffusion of macromolecules within the cytoplasm. We measured diffusion within living cells of different sizes by imaging and analyzing the diffusive motion of genetically-encoded multimeric cytoplasmic nanoparticles (cytGEMs), 40 nm-diameter fluorescent particles which inform on the diffusion of macromolecular complexes that are approximately the size of ribosomes (Delarue *et al*., 2018; Lemière *et al*., 2022; Molines *et al*., 2022). By analyzing cell size mutant strains, we found that diffusion within the cytoplasm decreased in a small cell mutant and increased in large cell mutants. Analyses of cytokinesis mutants demonstrated that this change in large cells was dependent on the DNA-to-Cytoplasm ratio. To gain mechanistic insight into these cell size-dependent effects, we discovered changes in the composition of the proteome and notably, in ribosomal concentration, which explains at least some of these effects on diffusion (Delarue *et al*., 2018). Our studies reveal that cell size impacts the physical properties of the cytoplasm, providing new perspectives into how cell size affects cellular physiology and function.

## RESULTS

### Nanoparticle diffusion in the cytoplasm increases with cell size

To investigate the relationship between cell size and intracellular diffusion, we expressed and imaged 40nm cytGEMs nanoparticles in *S. pombe* wildtype and cell size mutant cells, and through analyses of their motion, determined the effective diffusion coefficient in each strain (Delarue *et al*., 2018; Lemière *et al*., 2022; Molines *et al*., 2022). We grew *wee1-50*, wildtype, and *cdc25-22* cells at the permissive temperature 25°C and then shifted the cultures to the non-permissive temperature of 36°C for 6 hr before imaging (Fig. 1 A). At this temperature, *wee1-50* cells exhibit cell cycles with shorter G2 phases and enter mitosis at an abnormally short length, while *cdc25-22* cells arrest in G2 phase and continue to grow in length (Nurse, 1975; Fantes and Nurse, 1978). As these rod-shaped cells maintain approximately similar cell widths, the length of the cell was used as a proxy of cellular volume (Mitchison, 1957; Facchetti *et al*., 2019; Knapp *et al*., 2019). Upon the temperature shift, we showed that *wee1-50*, wildtype, and *cdc25-22* cells exhibited an average cell length of 5.61 ± 0.3 µm, 10.84 ± 1.39 µm, and 38.32 ± 1.36 µm, respectively (mean ± STD of replicate experiments). Measurement of the effective diffusion coefficient of cytGEMs in each cell population yielded average cytGEMs effective diffusion coefficients of 0.41 ± 0.04 µm^2^/s in *wee1-50*, 0.63 ± 0.07 µm^2^/s in wildtype, and 0.86 ± 0.04 µm^2^/s in *cdc25-22* cells (Fig. 1 B-C; mean ± STD of replicate experiments). Thus, cytGEMs diffusion showed a striking positive correlation between cell size and nanoparticle diffusion in the cytoplasm at the population level. We then analyzed the relationship between cytGEMs diffusion and cell size in individual cells, which exhibited a similar trend of increasing diffusion with cell size (Fig. S1 D). Positive correlations between cell length and cytGEMs diffusion (using a simple linear regression weighted by number of trajectories per cell) were also apparent when analyzing individual cells within *wee1-50* and *cdc25-22* strains (Fig. 1 D, Fig. S1 A-C). However, no significant changes were seen in range of sizes exhibited in wildtype cells, as noted previously (Fig. 1 D; Fig. S1 B) (Garner *et al*., 2023).

**Figure 1.**
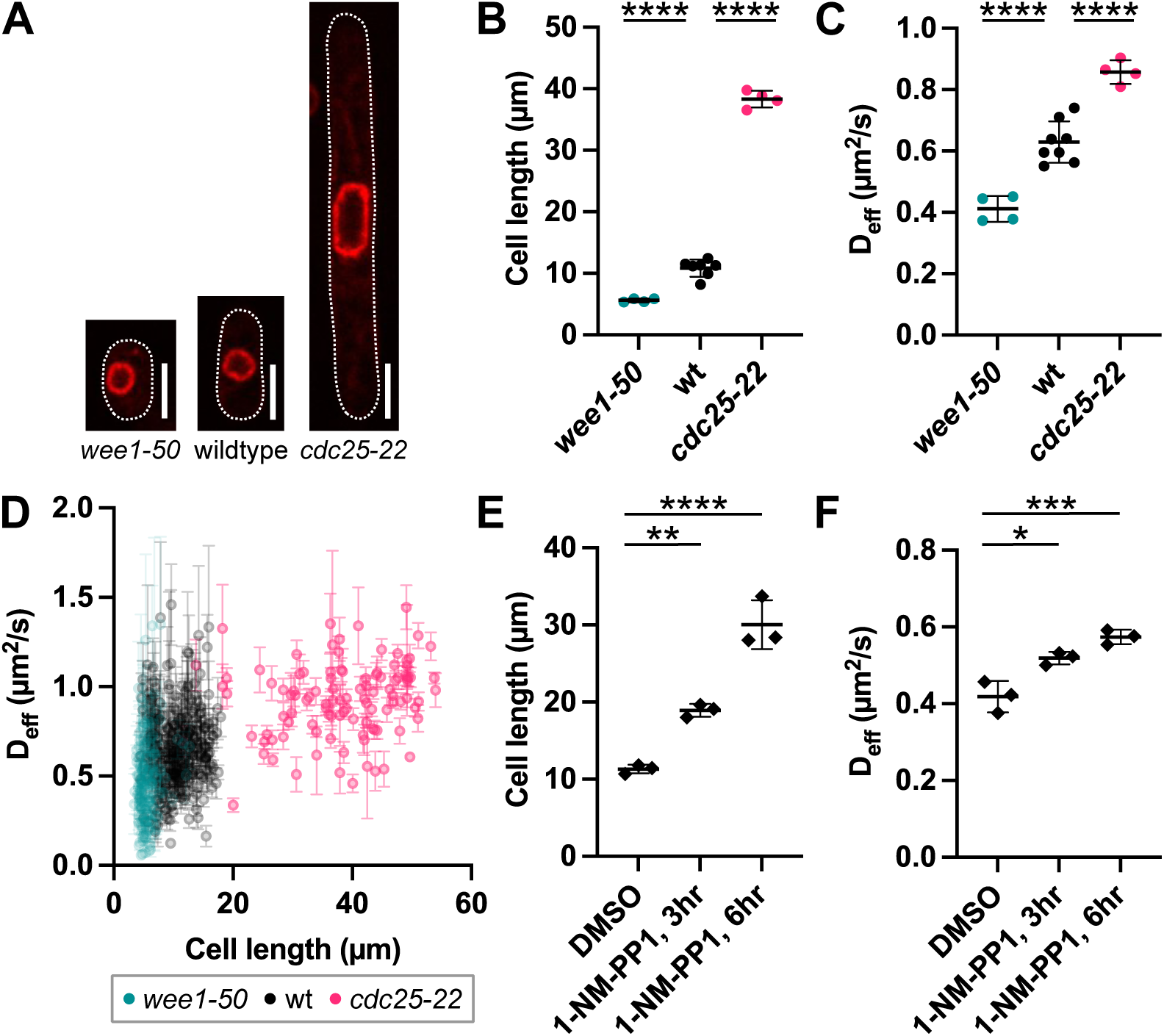
Nanoparticle diffusion increases with cell size. (**A**) Images (sum projection of 3 middle slices) of *wee1-50*, wildtype, and *cdc25-22* cells with nuclear membrane marker Ish1-GFP (red) grown at the permissive temperature 25°C overnight and shifted to the non-permissive temperature 36°C for 6 hr before imaging. Scale bar is 5μm. (**B**) Cell length (mean ± STD of replicate experiments; N_CELLS_ ≥ 106 per condition from at least 4 biological replicates) (1-way ANOVA, p < 0.0001) and (**C**) cytGEMs diffusion coefficients (mean ± STD of replicate experiments; N_GEMS_ ≥ 3183 per condition from at least 4 biological replicates) for *wee1-50*, wildtype, and *cdc25-22* cells grown with the temperature shift protocol described in (**A**) (1-way ANOVA, p < 0.0001). (**D**) Cell length and cytGEMs diffusion coefficients (mean ± SEM of cytGEMs trajectories per cell; N_CELLS_ ≥ 106 per condition from at least 4 biological replicates) plotted for individual cells for *wee1-50*, wildtype, and *cdc25-22* cells grown with the temperature shift protocol described in (**A**). (**E**) Cell length (mean ± STD of replicate experiments; N_CELLS_ ≥ 113 per condition from 3 biological replicates) and (**F**) cytGEMs diffusion coefficients (mean ± STD of replicate experiments; N_GEMS_ ≥ 5709 per condition from 3 biological replicates) for *cdc2-asM17* cells treated with 0.25% DMSO or 10μM ATP analog 1-NM-PP1. (1-way ANOVA, * - p < 0.05, ** - p < 0.01, *** - p < 0.001, **** - p < 0.0001).

To address concerns of possible effects of temperature shifts on diffusion and cytoplasmic viscosity, we grew *wee1-50*, wildtype, and *cdc25-22* cells at semi-permissive temperatures of 25°C and 28°C in steady state conditions. At these temperatures, the mutants generally exhibited significant, but more modest changes in cell size compared to those in the cells shifted to 36°C. While cytGEMs diffusion coefficients for each strain decreased with temperature, we still observed modest positive trends with cell size across the three strains at the population level (Fig. S1 D-G).

To generate large G2-arrested cells using another approach, we inhibited the analog-sensitive CDK1 allele *cdc2-asM17* with the ATP analog 1-NM-PP1 at 30°C (Aoi *et al*., 2014). These cells arrest in G2-phase and grow into large mononucleate cells, much like the *cdc25* mutants (Fig. 1 E). While untreated *cdc2-asM17* cells had an average cell length of 11.32 ± 0.57 µm, *cdc2-asM17* cells treated with 1-NM-PP1 for 3 hr and 6 hr had average cell lengths of 18.93 ± 0.84 µm and 30.05 ± 3.18 µm, respectively (mean ± STD of replicate experiments). The average cytGEMs diffusion coefficient was 0.42 ± 0.04 µm^2^/s for control cells, 0.52 ± 0.02 µm^2^/s for cells treated with 3 hr of 1-NM-PP1, and 0.57 ± 0.02 for cells treated with 6 hr of 1-NM-PP1 (mean ± STD of replicate experiments) (Fig. 1 F). Control treatments did not alter cytGEMs diffusion (Fig. S1 G). Overall, these results from a variety of strains and conditions showed a striking positive correlation of intracellular diffusion with cell size.

**Supplementary Figure 1.**
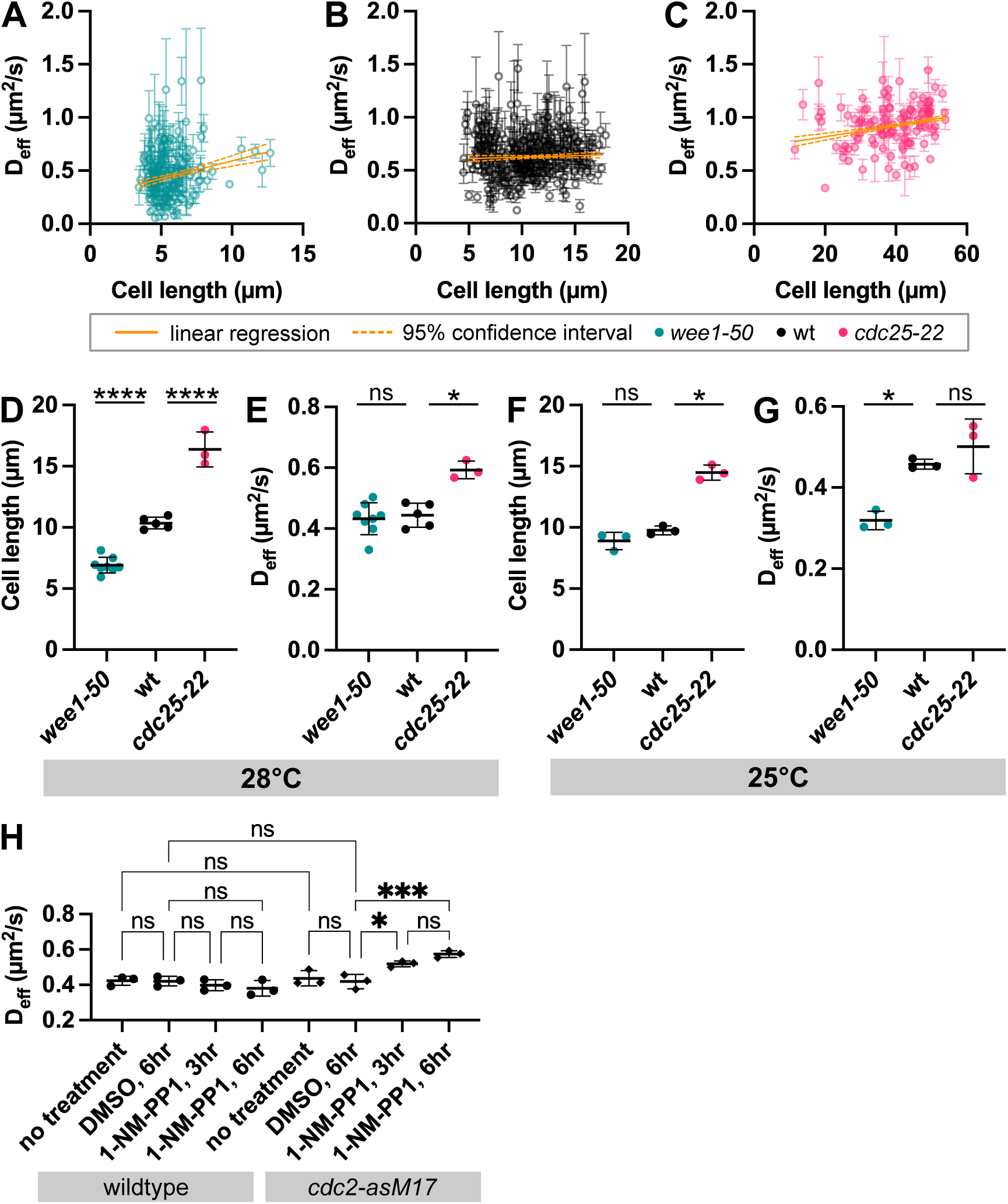
Intracellular diffusion increases with cell size in cell size mutant strains and at different temperatures. (**A**) Cell length and cytGEMs diffusion coefficients (mean ± SEM of cytGEMs trajectories per cell) plotted for individual cells for *wee1-50*, (**B**) wildtype, and (**C**) *cdc25-22* cells grown with the temperature shift protocol described in Figure 1A. Weighted linear regression (orange solid line) with 95% confidence interval (orange dashed lines) shown. Best-fit slopes are 0.03, 0.003, and 0.005 for (**A**), (**B**), and (**C**), respectively. (**D**) Cell length and (**E**) cytGEMs diffusion coefficients (mean ± STD of replicate experiments; N_GEMS_ ≥ 5135 per condition from at least 3 biological replicates) for *wee1-50*, wildtype, and *cdc25-22* cells grown at the steady-state semi-permissive temperature 28°C. (**F**) Cell length and (**G**) cytGEMs diffusion coefficients (mean ± STD of replicate experiments; N_GEMS_ ≥ 5981 per condition from at least 3 biological replicates) for *wee1-50*, wildtype, and *cdc25-22* cells grown at the steady-state permissive temperature 25°C. (**H**) CytGEMs diffusion coefficients (mean ± STD of replicate experiments) in wildtype and *cdc2-asM17* cells with varying conditions of 0.25% DMSO and 10μM 1-NM-PP1. (1-way ANOVA, * - p < 0.05, *** - p < 0.001, **** - p < 0.0001).

### Nanoparticle diffusion is maintained in large multinucleate cells

To determine whether the increase in cytGEMs diffusion in larger cell sizes was due to a decrease in the DNA-to-Cytoplasm ratio, we analyzed large multinucleated fission yeast cells in which the DNA-to-Cytoplasm ratio is not changed. We generated these multinucleate cells using the well-established mutants *sid2-as* and *cdc11-119* that are defective in the SIN regulatory pathway of cytokinesis (Nurse *et al*., 1976; Grallert *et al*., 2012). These conditional mutants continue to grow in length and undergo nuclear division cycles in the absence of septation. *sid2-as* cells treated with the ATP analog 1-NM-PP1 formed progressively larger cells with multiple nuclei at 3 and 6 hr of treatment (Fig. 2 A-C). Control *sid2-as* cells had an average cell length of 10.36 ± 0.34 µm, while *sid2-as* cells treated with 1-NM-PP1 for 3 hr and 6 hr had average cell lengths of 16.55 ± 0.91 µm and 25.95 ± 1.6 µm, respectively (mean ± STD of replicate experiments). Based on the average cell length and number of nuclei per condition, we estimated that the DC ratio of 1-NM-PP1-treated cells did not decrease compared to the control. Despite being larger in cell size, we found that cytGEMs diffusion coefficients in treated *sid2-as* cells (3 hr: 0.48 ± 0.6 µm^2^/s; 6 hr: 0.56 ± 0.01 µm^2^/s) were comparable with the control cells (0.52 ± 0.04 µm^2^/s) (mean ± STD of replicate experiments) (Fig. 2D, S2A).

**Figure 2.**
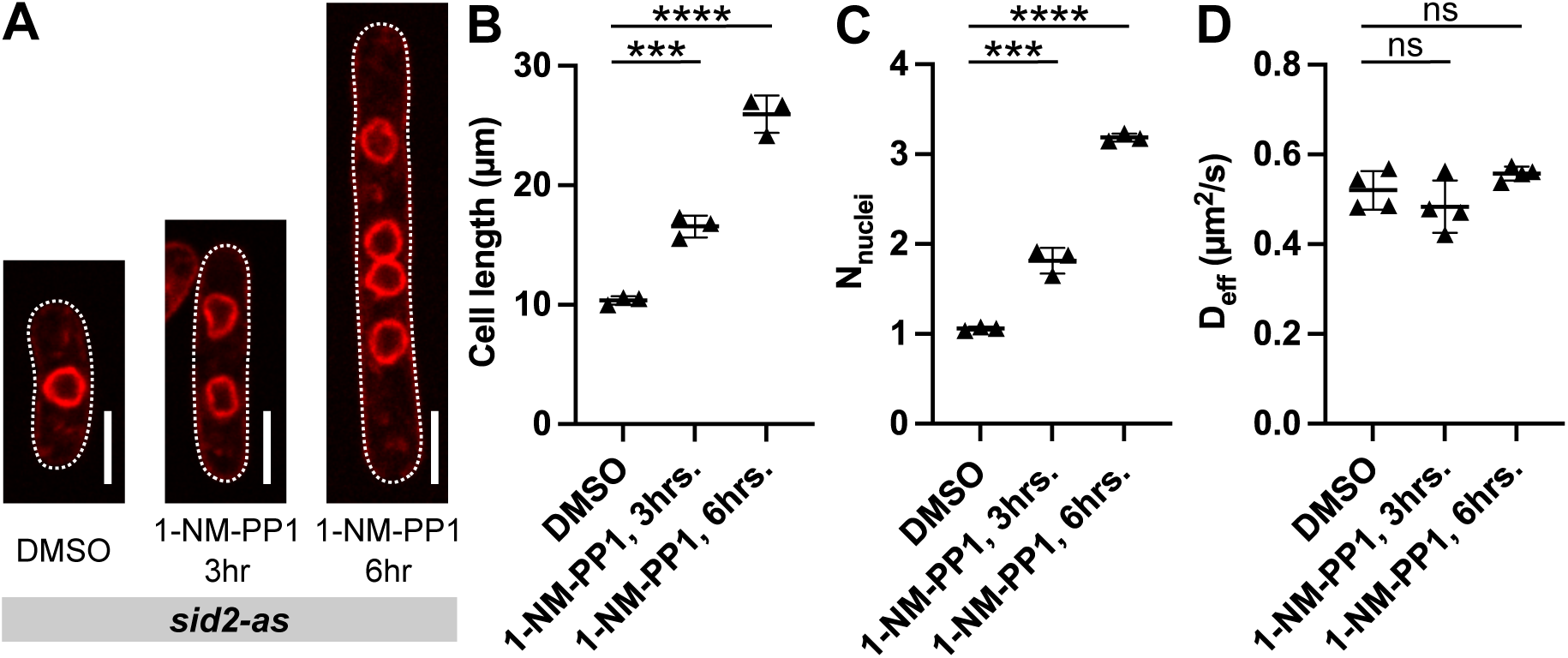
Nanoparticle diffusion does not change with cell size in large multinucleate cells. (**A**) Images (sum projection of 3 middle slices) of *S. pombe sid2-as* cells with nuclear membrane marker Ish1-mScarlet (red). Cells were grown at steady-state 30°C and treated with 0.25% DMSO or 10μM ATP analog 1-NM-PP1. Left to right: 6hr DMSO, 3hr 1-NM-PP1, and 6hr 1-NM-PP1. Scale bar is 5μm. (**B**) Cell length, (**C**) number of nuclei (mean ± STD of replicate experiments; N_CELLS_ ≥ 140 per condition from 3 biological replicates), and (**D**) cytGEMs diffusion coefficients (mean ± STD of replicate experiments; N_GEMS_ ≥ 5546 per condition from 4 biological replicates) for *sid2-as* cells described in (**A**). (1-way ANOVA, *** - p < 0.001, **** - p < 0.0001).

Next, we inhibited cytokinesis by using the temperature-sensitive mutant *cdc11-119* (Nurse *et al*., 1976). Wildtype cells and *cdc11-119* cells were grown at the permissive temperature 25°C overnight and shifted to the non-permissive temperature 36°C for 3 hr. We observed comparable cytGEMs diffusion coefficients in the *cdc11-119* cells compared to control populations (Fig. S2 B). Overall, these results suggest that a decrease in DNA-to-Cytoplasm ratio rather than an increase in cell size alone, underlies the increase in intracellular diffusion observed in large cells (Fig. 1).

**Supplementary Figure 2.**
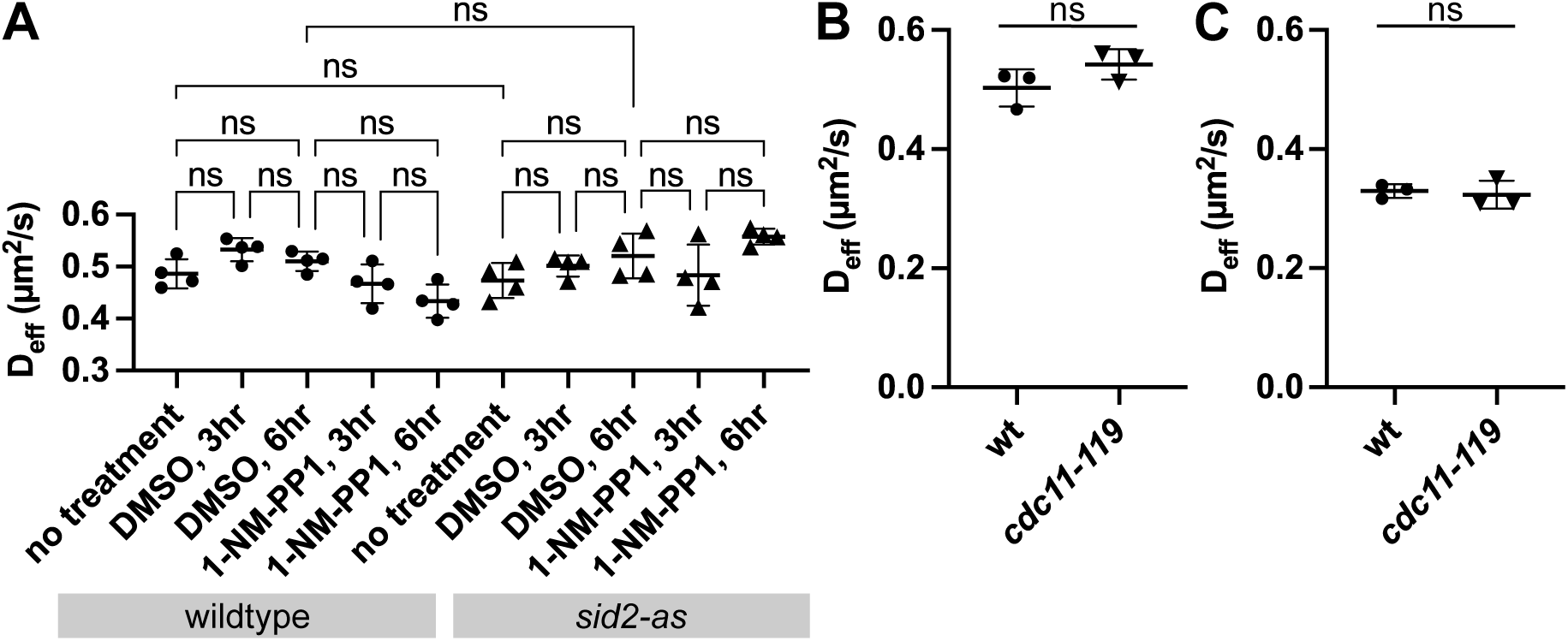
Diffusion coefficients are comparable between large multinucleate cells and wildtype cells. (**A**) CytGEMs diffusion coefficients (mean ± STD of replicate experiments) in wildtype and *sid2-as* cells with varying conditions of 0.25% DMSO and 10μM 1-NM-PP1 (1-way ANOVA, p > 0.05). (**B**) CytGEMs diffusion coefficients (mean ± STD of replicate experiments; N_GEMS_ ≥ 7630 per condition from 3 biological replicates) for wildtype and *cdc11-119* cells grown at permissive temperature 25°C overnight and shifted to the non-permissive temperature 36°C for 3 hr before imaging (Komogorv-Smirnov test, p = 0.6). (C) CytGEMs diffusion coefficients (mean ± STD of replicate experiments; N_GEMS_ ≥ 9782 per condition from 3 biological replicates) for wildtype and *cdc11-119* cells grown at the steady-state permissive temperature 25°C (Komogorv-Smirnov test, p = 0.6).

### Ribosomal and total protein concentrations decrease in large cells

We hypothesized that the cell size-dependent changes in cytGEMs diffusion reflect changes in cytoplasmic composition or concentration. Previous studies in other organisms suggest that an increase in diffusion can correlate with a decrease in ribosome concentration or overall protein concentration (Delarue *et al*., 2018; Neurohr *et al*., 2019; Molines *et al*., 2022).To assess ribosomal concentration, we measured the fluorescence intensity of Rps2-GFP, a functional fusion of the essential small subunit ribosomal protein expressed at the native locus (Knapp *et al*., 2019; Lemière *et al*., 2022). *wee1-50*, wildtype, and *cdc25-22* cells expressing Rps2-GFP were grown at the permissive temperature 25°C overnight and shifted to the non-permissive temperature 36°C for 6 hr before imaging (Fig. 3 A). To facilitate equivalent processing, cells of the three strains were mixed and imaged in the same field. We found a distinct inverse relationship of Rps2 intensity with cell size (Fig. 3 B). In binned data, the average intensity of Rps2-GFP was significantly lower in bigger cells (cell length ≥ 18 µm) compared to medium-sized cells (cell length between 9 and 18 µm), but no significant differences were detected between medium and smaller cells (cell length ≤ 9 µm) (Fig. 3 B-C).

**Figure 3.**
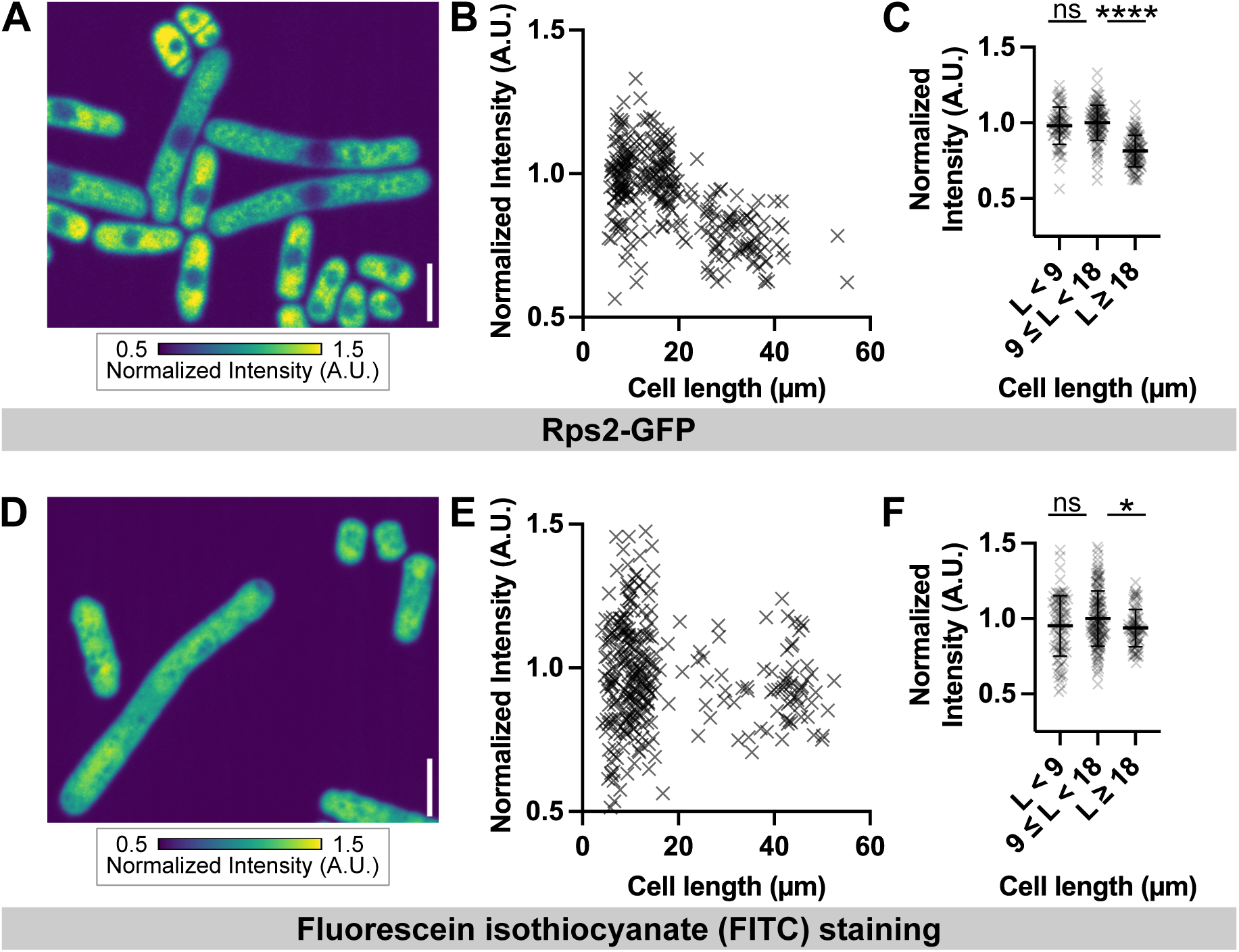
Large cells have decreased concentrations of ribosomes. (**A**)Image (sum projection of 3 middle slices) of a mixture of *wee1-50*, wildtype, and *cdc25-22* live cells with ribosomal protein marker Rps2-GFP. Scale bar is 5μm. (**B**) Rps2-GFP intensity and length per cell. **(C)** Rps2-GFP intensities (mean ± STD per cell length category; N_CELLS_ ≥ 72 per condition from 3 biological replicates) measured in a mixture of *wee1-50*, wildtype, and *cdc25-22* cells and categorized by cell length. (**D**) Image (sum projection of 3 middle Z-slices) of a mixture of *wee1-50*, wildtype, and *cdc25-22* fixed cells, treated with RNase A, and stained with FITC. (**E**) FITC intensity and length per cell. (**F**) FITC intensities (mean ± STD per length category; N_CELLS_ ≥ 73 per condition from 3 biological replicates) measured in a mixture of *wee1-50*, wildtype, and *cdc25-22* fixed cells and categorized by cell length. Intensity values for Rps2-GFP and FITC are normalized to the mean intensity of the 9 ≤ L<18 category (1-way ANOVA test, * - p < 0.05, **** - p < 0.0001).

To assess overall protein concentration, we measured the intensity of fluorescent dye fluorescein isothiocyanate (FITC) staining (Fig. 3D) (Knapp *et al*., 2019). *Wee1-50*, wildtype, and *cdc25-22* cells were shifted to 36°C for 6 hr, mixed, fixed, stained with FITC and imaged. In data binned by cell size, compared to medium-sized cells, mean FITC intensity was about 6% lower in bigger cells (1-way ANOVA, p=0.04) and 5% lower in smaller cells (1-way ANOVA, p=0.1). These results were consistent with an overall decrease in dry mass density seen previously in *cdc25-25* cells (Odermatt *et al*., 2021).

Overall, our results showed that larger cells exhibited a decrease in ribosomal protein concentration and to a lesser extent, overall protein concentration, which begin to provide an explanation for the increase in cytGEMs mobility with increasing cell size.

### Proteome composition varies with cell size

Finally, to gain further insight into the mechanisms underlying changes in diffusion, we investigated how the molecular composition of the cytoplasm changes with cell size by comparing the proteomes of *S. pombe wee1-50*, wildtype, and *cdc25-22* cells. Mass spectrometry was analyzed using SILAC in pairwise comparisons. First, *wee1-50* and *cdc25-22* SILAC strains were both labeled with varying lysine and arginine isotopes at the permissive temperature 25°C overnight and shifted to the non-permissive temperature 36°C for 6.5 hr, which produced similar size ranges to the *wee1-50* and *cdc25-22* cell populations analyzed in Figure 1. Proteomic analyses detected 3,353 proteins out of 5,117 identified *S. pombe* proteins (∼65% coverage) (The UniProt Consortium, 2022), and our two experimental repetitions yielded consistent results (R=0.83, Pearson) (Fig. 4 A). We categorized proteins by their subcellular location or macromolecular complex such as histones (magenta), ribosomes (orange), and ER (cyan) (Fig. 4 A). Finally, we grouped proteins by their subcellar location or macromolecular complex and averaged their collective ratios (Fig. 4 B). Because relative concentrations of each protein were calculated and normalized within each strain, we note that these analyses cannot reveal alterations in real protein concentrations but only relative changes to other proteins.

**Figure 4.**
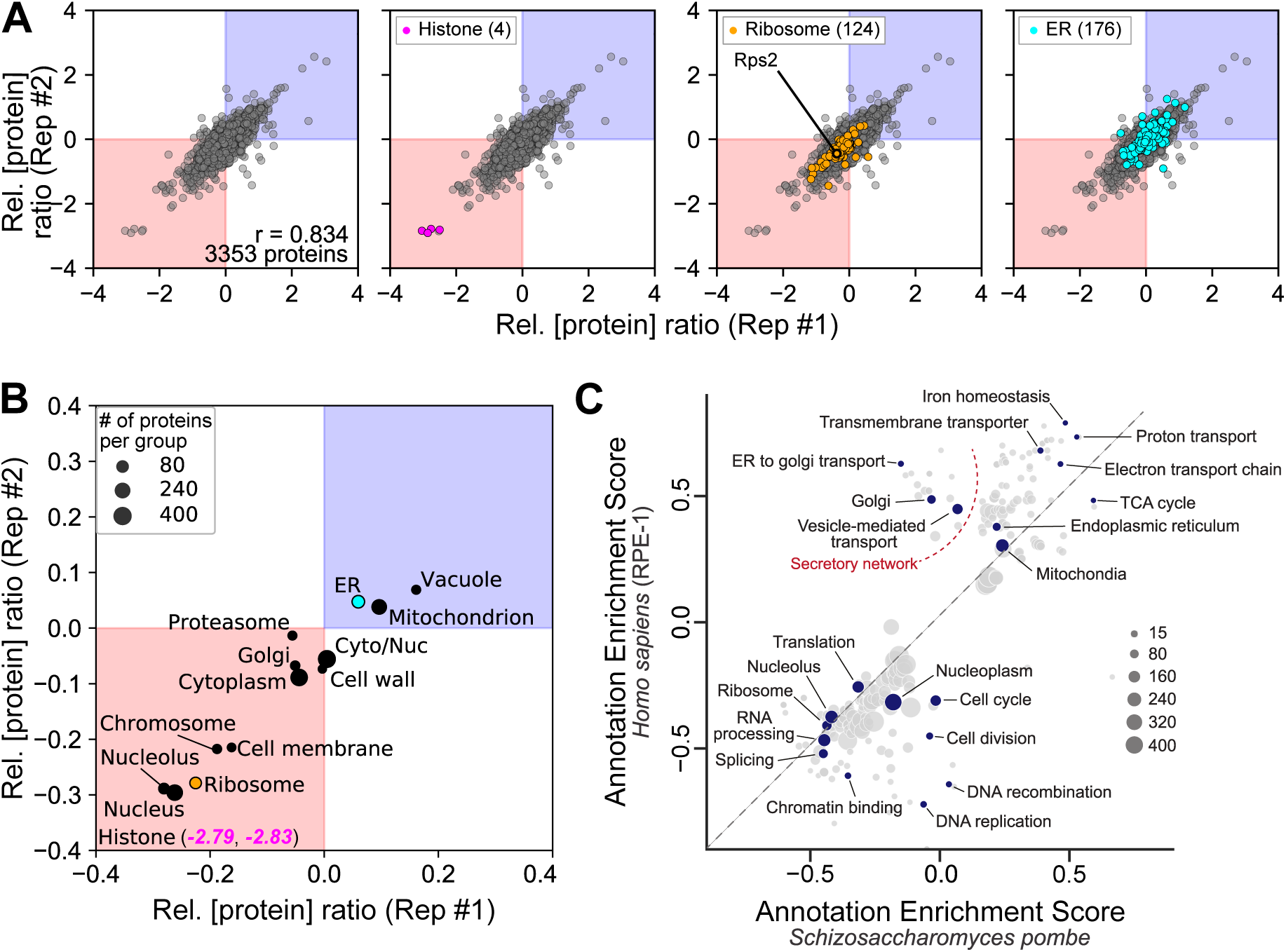
Proteome composition varies with cell size. The proteomes of *cdc25-22* and *wee1-50* cells grown at 25°C and shifted to 36°C for 6.5 hr were compared using SILAC mass spectrometry. Concentrations of each protein were determined per strain and normalized to the respective strain’s proteome. (**A**) Relative protein concentration ratios (*cdc25-22*/*wee1-50*) for each detected protein; ratios from two replicates are plotted. Upper right quadrant (blue) indicates superscaling proteins that have relative protein concentration ratios that are more than 1. These proteins are relatively more abundant in *cdc25-22* compared to *wee1-50*. By contrast, the lower left quadrant (red) indicates subscaling proteins that have relative protein concentration ratios that are less than 1. These proteins are relatively less abundant in *cdc25-22* compared to *wee1-50*. Histone proteins, ribosomal proteins and ER proteins (with the number of proteins represented in parentheses) are highlighted to demonstrate different scaling relationships at different subcellular locations. (**B**) Average relative concentration ratios of proteins grouped by subcellular localization in comparison of two replicates. (**C**) 2D annotation enrichment analysis using the Protein Slope values paired by sequence ontology between fission yeast and cultured human cells from Lanz *et al*. 2022. The identity line is dashed. Each dot is an annotation group, and the position of the dot is determined by the mean slope value of the proteins in the group (rank-based). Positive and negative enrichment scores indicate groups of super-scaling and sub-scaling proteins, respectively.

Overall, we observed differential scaling of proteins when comparing the proteomes of large and small cells. Proteins associated with the nucleus including the nucleolus, histones (magenta, out of range), and chromosome sub-scaled with cell size, i.e. they were underrepresented in the large *cdc25-22* cells compared to small *wee1-50* cells (Fig. 4 B, red quadrant). This sub-scaling behavior was expected, as chromosome-associated proteins such as histones are known to scale with DNA content, not cell size (Amodeo *et al*., 2015; Claude *et al*., 2021). Notably, ribosomal proteins (Fig. 4 B, orange) also exhibited sub-scaling, which supported our observation that ribosome concentration is decreased in these large cells (Fig. 4 A-C). Overall cytoplasmic proteins are also sub-scaling with cell size. In contrast, proteins associated with the endoplasmic reticulum (cyan), mitochondria, and vacuoles super-scaled with cell size, i.e. they were overrepresented in the large *cdc25-22* cells compared to small *wee1-50* cells (Fig. 4 B, blue quadrant).

Next, we examined the proteome data for scaling of cellular processes and signaling pathways implicated in regulation of cytoplasmic properties in other studies. One candidate signaling pathway that regulates ribosome concentration and stress responses is the TORC1 pathway (Delarue *et al*., 2018). Proteins associated with TORC complexes and ribosome biogenesis sub-scaled with cell size (Fig. S3 A). To test whether TORC1 activity was decreased in large cells, we found that factors downstream of the TORC1-regulated Sfp1 transcription factor also sub-scaled with size (Tai *et al*., 2023). However, in contrast to other studies characterizing cell size proteome changes (Neurohr *et al*., 2019; Lanz *et al*., 2022, 2024), we detected no significant super-scaling effects on stress-associated pathways such as the core environmental stress response (CESR) (Chen *et al*., 2003). As small viscogens such as trehalose and glycerol may be additional factors in cytoplasmic viscosity, we found that proteins involved in trehalose biosynthesis sub-scaled with size and those associated with trehalose breakdown super-scaled with size (Fig. S3 A) (Sakai *et al*., 2024). To analyze the top hits for sub- and super-scaling proteins in our data set, we performed a gene ontology enrichment analysis (PANTHER overrepresentation test) (Fig. S3 B). Top super-scaling proteins were generally involved in metabolic pathways associated with membrane-bound organelles, whereas top sub-scaling proteins were associated with cell polarity regulation at cell tips, and mRNA regulation and gene expression in the nucleus. Of interest, among the sub-scaling cell polarity proteins was the DYRK protein kinase Pom1 as well its regulators Tea1 and Tea4, which all localize to cell tips and contribute to cell size sensing for cell size regulation (Martin and Berthelot-Grosjean, 2009; Moseley *et al*., 2009; Hachet *et al*., 2011; Wood and Nurse, 2015).

Additional proteomic comparisons between *wee1-50*, wildtype, and *cdc25-22* cells grown under other conditions supported these results. First, ratios of the proteomes of *cdc25-22* and wildtype cells that were shifted to 36°C for 6.5 hr (as described in Fig. 4) showed similar trends as our comparison between *cdc25-22* and *wee1-50* cells (Fig. S4 A-B). Second, we compared *cdc25-22* and *wee1-50* strains grown at steady-state at 28°C (similar to Fig. S1 D-E) to remove effects of temperature shift. Here, we observe the same general trends in the proteome, with the notable exception of ribosome proteins which scaled with cell size in these conditions (Fig. S4 C-D).

Our data showed striking similarities with a proteome data set that compared small and large cell proteomes in mammalian cells. An annotation-based analyses comparing sub-scaling and super-scaling proteins in *H. sapiens* RPE-1 cells (Lanz *et al*., 2022, 2024) with our data on *cdc25-22/ wee1-50* in *S. pombe* (Fig. 4A, B) revealed notable conservation (Fig. 4C). For instance, the sub-scaling of ribosomal and chromatin-associated proteins and super-scaling of endoplasmic reticulum and mitochondria proteins with cell size were conserved. A comparison of cell size-dependent changes also revealed similar relationships in the proteomes of RPE-1 cells and *S. cerevisiae* (Lanz *et al*., 2022, 2024). Thus, the composition of the proteome exhibits characteristic changes that are likely to be general conserved signatures of cell size in eukaryotes.

**Supplementary Figure 3.**
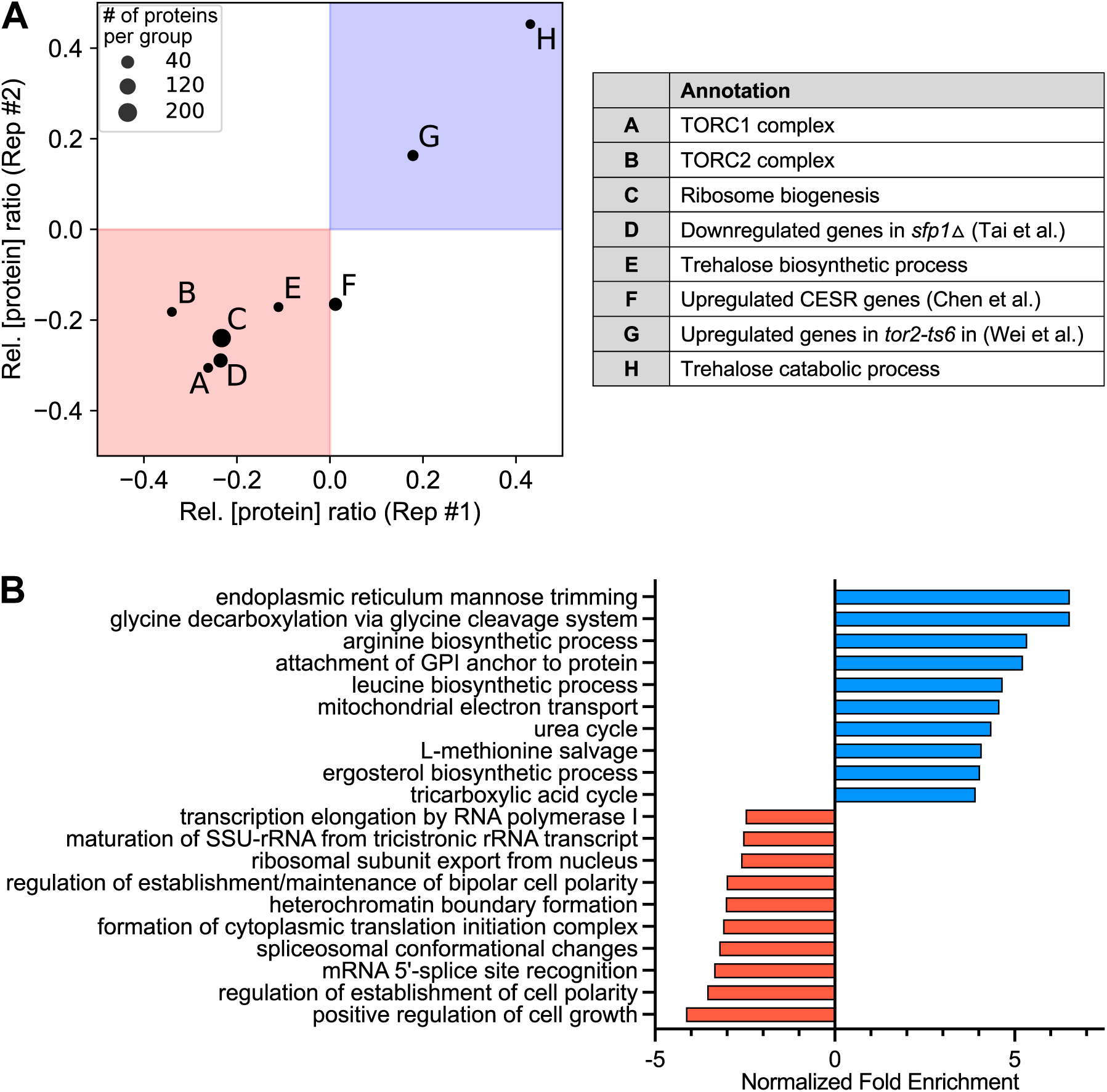
Comparison of *S. pombe* proteome size scaling with other studies. (**A**) Effect of cell size on scaling of proteins implicated in ribosomal regulation, TORC1 pathways, stress pathway and trehalose regulation. Relative concentration of proteins in *S. pombe cdc25-22* and *wee1-50* strains grown with 6.5 hr shift to 36°C as described in Figures 1 and 3. Ratios depict ratios of *cdc25-22*/*wee1-50*. (**B**) Gene ontology analysis of sub-(red) and super-scaling (blue) proteins in *cdc25-22*/*wee1-50* as above.

**Supplementary Figure 4.**
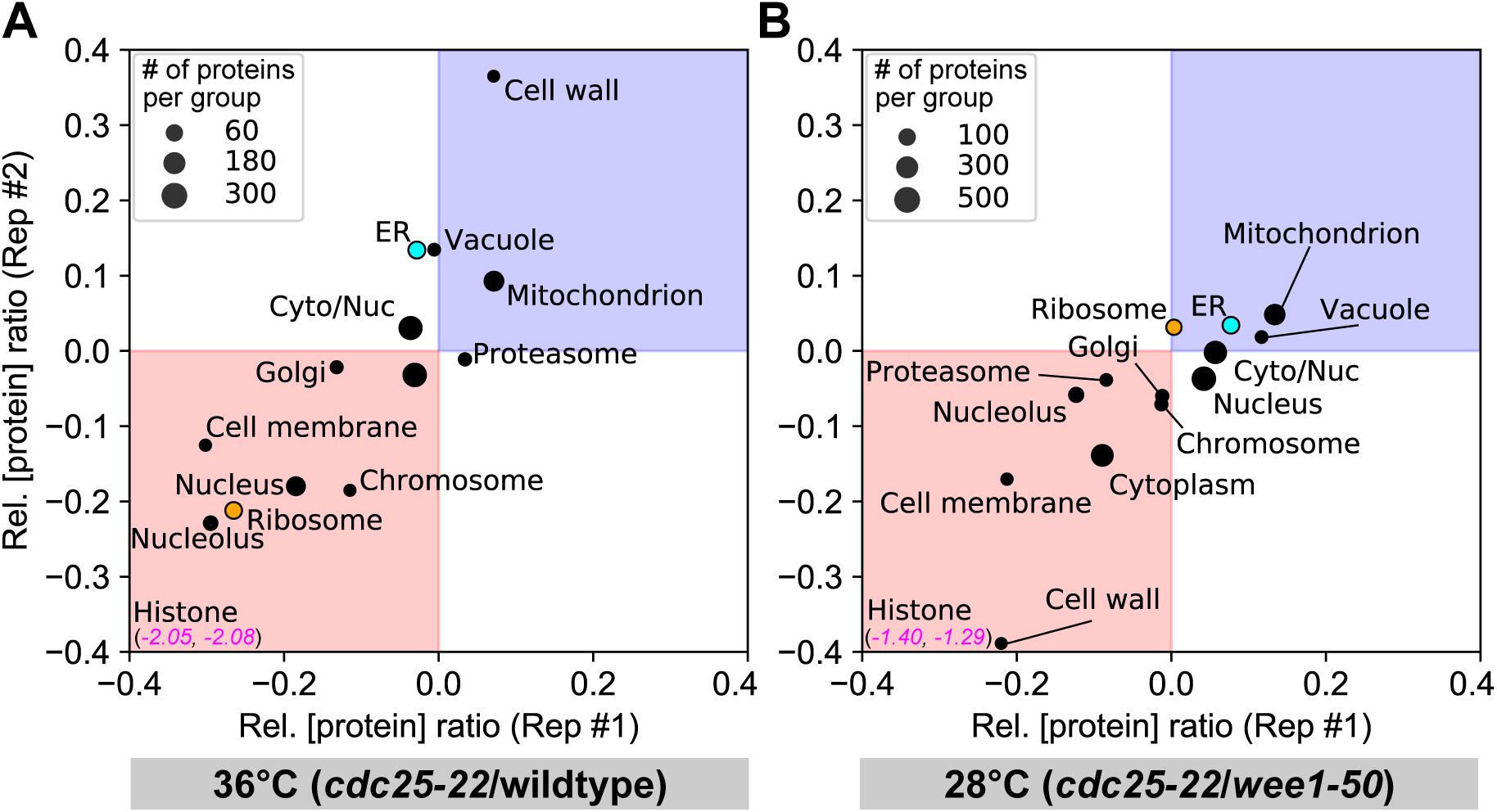
Cell size proteome changes at smaller cell size differences. Similar size-dependent effects on the proteome were seen in comparisons of fission yeast cells grown in other conditions. (**A**) *cdc25-22* and wildtype cells were grown at the permissive temperature 25°C and shifted to 36°C for 6.5 hr before sample collection. Plot shows the average relative protein concentration ratios (*cdc25-22*/wildtype) of proteins were grouped by subcellular localization in two replicates. (**B**) *cdc25-22* and *wee1-50* cells were grown at semi-permissive temperature of 28°C at steady state. Average relative concentration ratios (*cdc25-22*/*wee1-50*) proteins were grouped by subcellular localization in two replicates.

## DISCUSSION

Here we show that intracellular diffusion coefficients of macromolecular complex-sized particles exhibit a significant positive correlation with increasing cell size. Relative to wildtype cells, cytGEMs diffusion decreased in small *wee1-50* mutant cells and increased in large *cdc25-22* mutant cells (Fig. 1). However, cytGEMs diffusion was not changed in large multinucleate cells, demonstrating that DNA-to-Cytoplasm ratio may be the critical parameter that underlies diffusion rates rather than cell size alone (Fig. 2). In investigating the mechanism underlying these changes in diffusion, we showed that small and large cells exhibited different proteome compositions, with large cells exhibiting decreased concentrations of ribosome and nuclear proteins relative to other elements of the proteome. These results are consistent with a model in which diffusion increases in larger cells due to a decrease in the concentration of ribosomes and changes to the concentrations of many other cytoplasmic components. In proliferating cells, the effect of the DNA-to-Cytoplasm ratio suggests that a limiting factor may be the number of gene copies needed to maintain gene expression to support the exponential growth of the cytoplasm (Zhurinsky *et al*., 2010; Marguerat and Bähler, 2012; Neurohr *et al*., 2019; Balachandra *et al*., 2022; Cadart and Heald, 2022; Xie *et al*., 2022).

Overall, our study supports the premise that the properties of the cytoplasm vary at different cell sizes. While previous studies focus on the apparent dilution of the cytoplasm and/or changes in the biochemical composition in large cells (Neurohr *et al*., 2019; Lanz *et al*., 2022), our findings show that cell size impacts diffusion and crowding in the cytoplasm. As diffusion and crowding have broad range effects on the inner workings of the cell, including the rates of most biochemical reactions, our findings introduce a critical component in our understanding of the effects of cell size on cellular physiology.

Our findings on diffusion are consistent with a quantitative phase imaging study in fission yeast that showed cell cycle-dependent fluctuations in intracellular dry mass density, as well as a steady drop in density in *cdc25-22* arrested cells (Odermatt *et al*., 2021). This work suggests that density fluctuations arise from fluctuations in growth rate accompanied by a constant mass biosynthesis. Thus, one mechanism for density dilution in *cdc25-22* arrested cells may be due to the prolonged time in G2 phase when the rate of volume growth slightly outpaces mass biosynthesis. By contrast, the density increase in *wee1-50* cells may be due to a higher proportion of mitotic and dividing cells in the *wee1-50* population (Hagan *et al*.; Rowley *et al*., 1992). Thus, one mechanism for these changes in diffusion may be cell cycle dependent changes in the coordination between biosynthesis and volume growth.

Our studies contribute to a growing body of evidence that the cytoplasm not only becomes more dilute with increasing cell size, but also that the composition of the cytoplasm remodels with cell size. Comparison of our results with recent data in human cells and budding yeast cells show that this remodeling of the proteome is largely conserved in these eukaryotic cells (Fig. 4C) (Lanz *et al*., 2022, 2024). For example, in these three organisms, our studies agree that sub-scaling proteins are enriched in nuclear proteins, while super-scaling proteins are enriched in ER and mitochondrial proteins as well as metabolic proteins (Fig. 4; Supp Figure X). Proteome remodeling in budding yeast is thought to be independent of metabolic state but holds similarities to cells in the starvation state and during environmental stress response (Lanz *et al*., 2024). However, we did not detect these similarities for fission yeast (Fig. S3 A). Our proteome and fluorescence intensity analyses (Fig. 3, S3) together showed that the concentration of ribosomal proteins subscales in larger fission yeast cells. As ribosomes have been suggested to be a significant crowding agent of macromolecules in the cytoplasm (Delarue *et al*., 2018), the decrease in not only ribosomal protein concentration, but also the decrease in ribosome biogenesis proteins and TOR complex proteins, altogether provide a possible mechanism for increased diffusion mediated through the TOR pathway.

In addition to ribosomal concentration, it is likely that other factors also contribute to cell size effects. Ribosomal protein concentration cannot readily account for all the diffusion data; for instance, we noted several cases where cells exhibited changes in diffusion but no detectable changes in ribosomal protein concentration -- for instance in *wee1-50* cells (Fig. 1, 3) or lack of sub-scaling of ribosomal proteins in *wee1-50* and *cdc25-22* cells grown at 28°C (Fig. S1, S4). It is possible that the methods were not sensitive enough to detect subtle changes. However, in addition, there are likely to be other factors that contribute to diffusion changes, such as small viscogens like trehalose and glycerol (Fig. S3) or other crowding agents.

Changes in macromolecular crowding and diffusion are predicted to have significant impacts on the biochemistry and mechanobiology inside the cell. For instance, these physical cytoplasmic properties not only affect rates of biochemical reactions, dynamics of molecular conformational changes, and protein expression, but they also impact organelle size and phase transitions that help to organize the cytoplasm (Rivas and Minton, 2016; Mitchison, 2019; Marshall, 2020). Our studies suggest that one reason why cell size is maintained in a homeostatic manner is to maintain the state of the cytoplasm. In abnormally large cells seen in senescence, aging and disease states, altered cytoplasmic properties may contribute to slower growth rates, abnormal cellular function, and cell death (Neurohr and Amon, 2020; Xie *et al*., 2022). Future studies promise to reveal how cell size affects the intracellular environment responsible for various cellular functions.

## METHODS AND MATERIALS

**Table 1.**
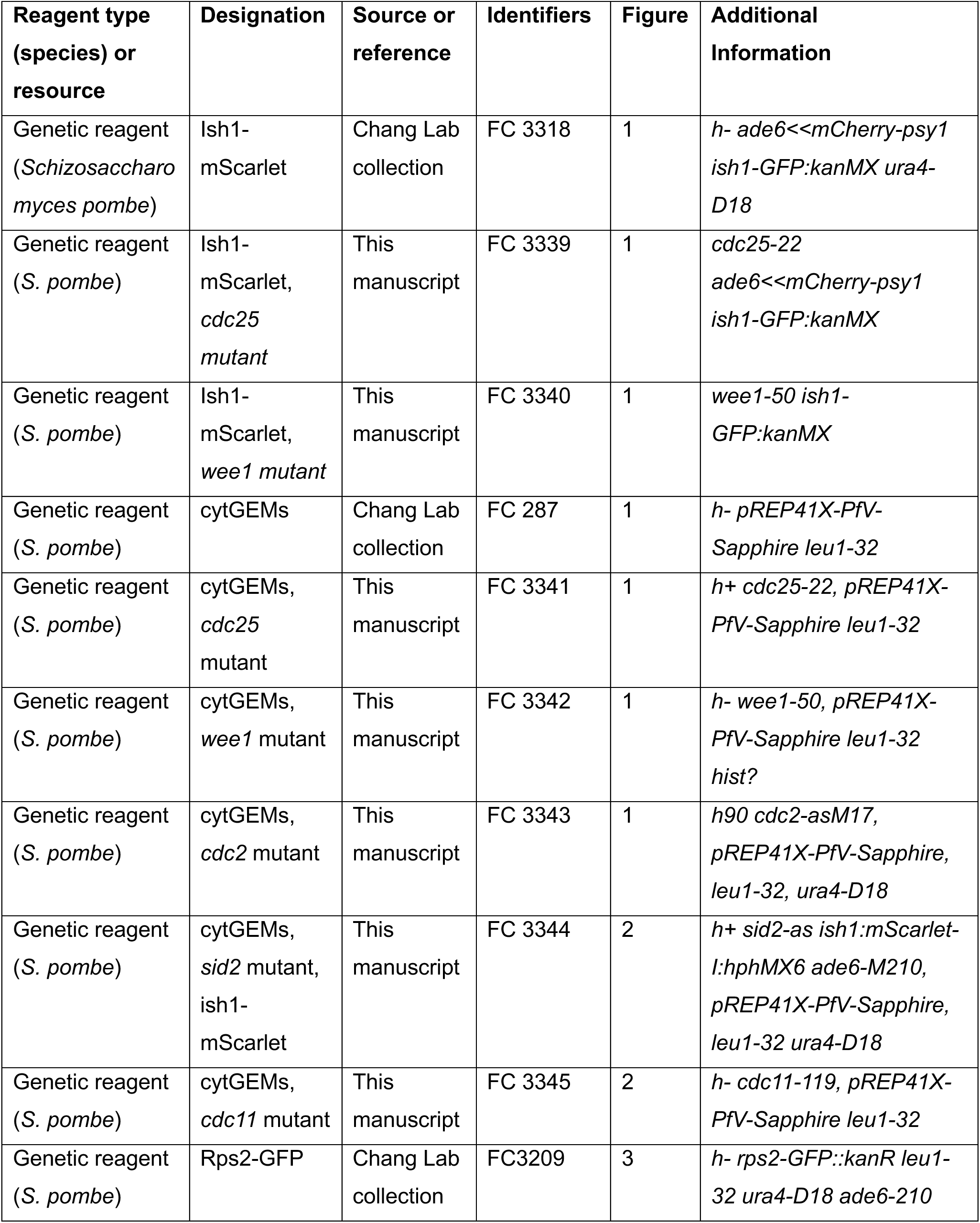
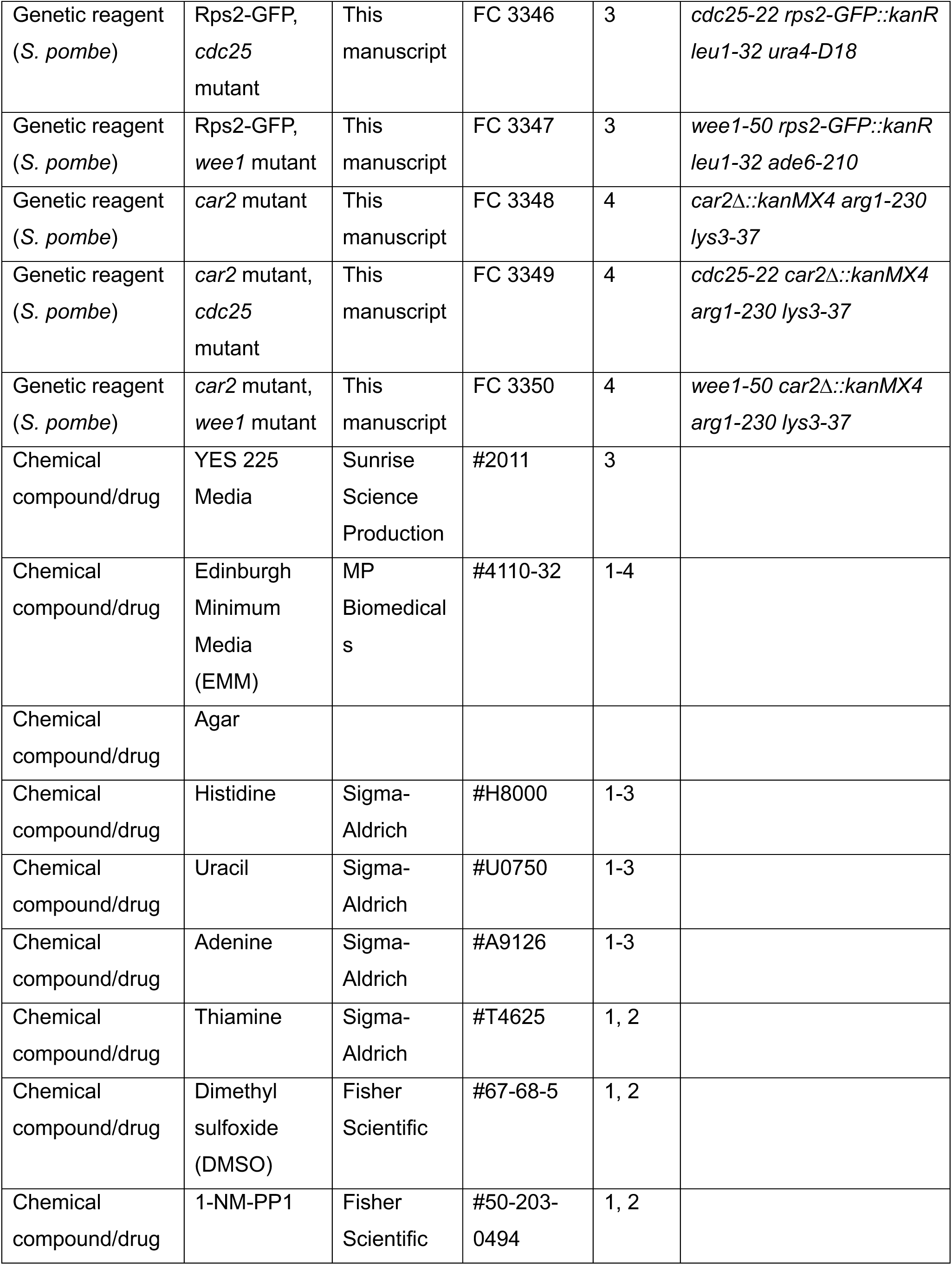
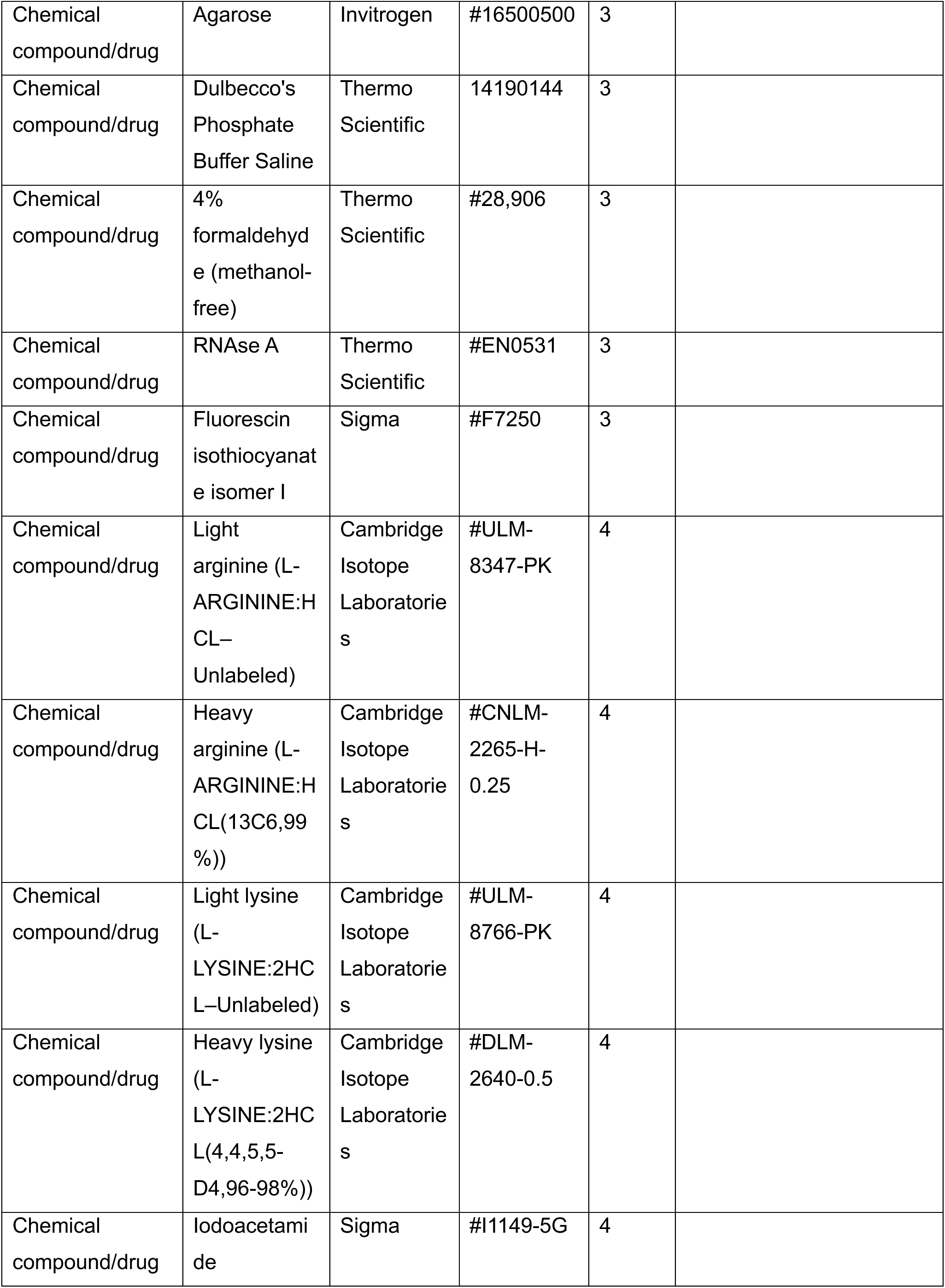
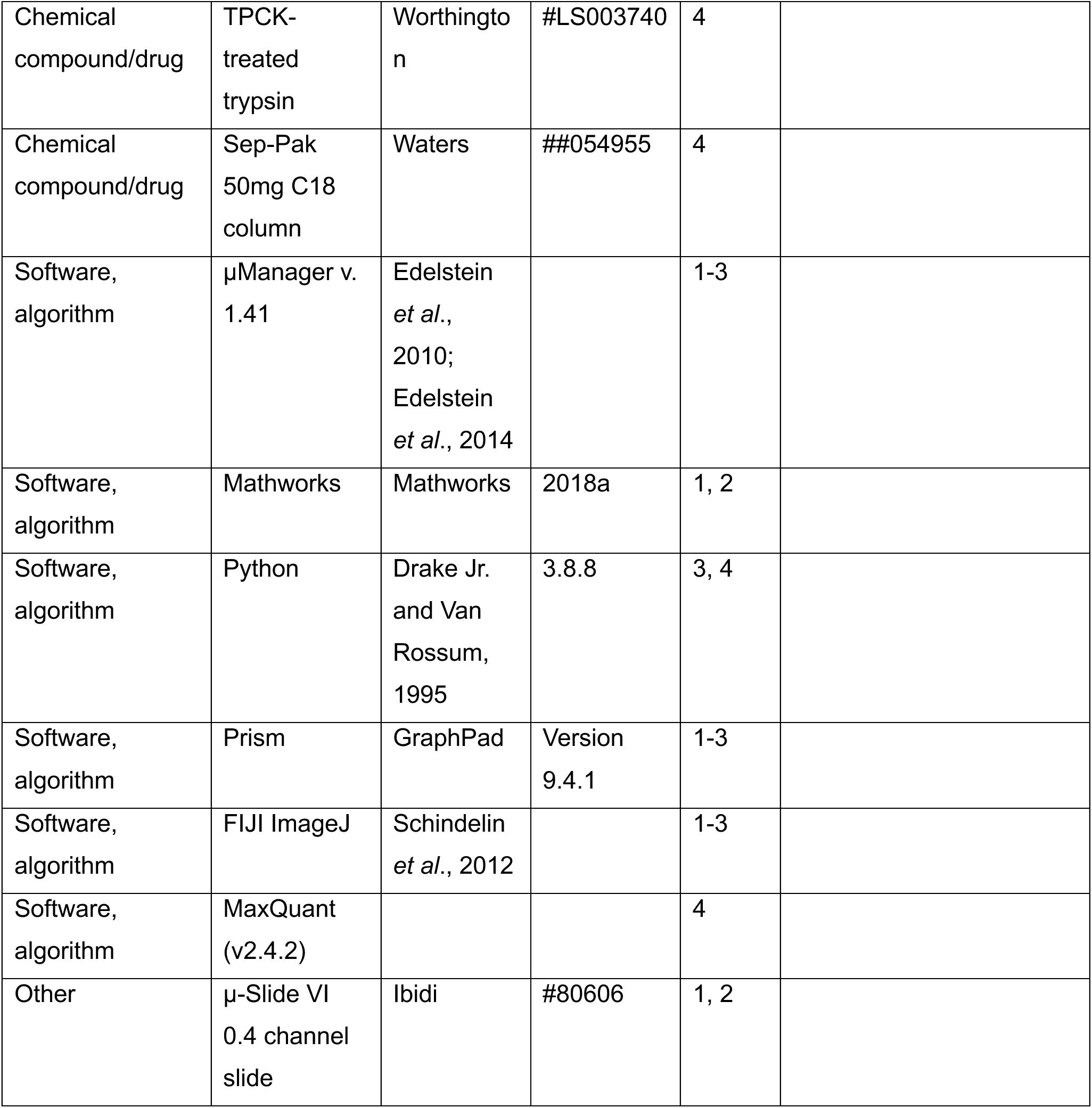
Key resources used in this study.

### Yeast strains and media

*Schizosaccharomyces pombe* strains were constructed and maintained using standard methods (Forsburg, 2003). The strains used in this study are listed in Table 4.1. For expression of 40nm cytGEMs, yeast cells were transformed with the plasmid pREP41X-PfV-mSapphire for expression of the protein fusion PfV encapsulin-mSapphire (Delarue *et al*., 2018; Lemière *et al*., 2022; Garner *et al*., 2023). These cells were grown in EMM3S (250 mg/mL adenine, 250 mg/mL histidine, 250 mg/mL uracil, and no leucine) media with 0.1 μg/mL thiamine for an intermediate level of expression from the nmt1* promoter to optimize the appropriate numbers of cytGEMs in each cell (Maundrell, 1993; Molines *et al*., 2022). In other experiments, cells were grown in rich YES (Fig. 3) or SILAC adjusted EMM media (Fig. 4) (Swaffer *et al*., 2016).

### Temperature shift and inhibitor treatments

Fission yeast cells of different cell sizes and ploidy were generated using established conditional cell cycle mutants (see main text). For temperature-shift experiments (Fig. 1 A-D, 2) wildtype and temperature-sensitive mutant cells were inoculated from colonies freshly grown from the frozen stocks on EMM3S (minus leucine) agar plates grown at 25°C for 3 days and stored at room temperature for less than 7 days. Cells were inoculated in liquid EMM3S medium and grown at 25°C with shaking for over 12 hr to exponential phase in the range of OD_600_ 0.2 to 0.6. The flasks were then transferred to a 36°C shaking incubator for the indicated period (3-6 hr). The cells were then harvested and mounted in chambers for imaging on the lab bench and promptly returned to 36°C in the pre-warmed microscope system incubator. No significant differences in cytGEMs diffusion were found when mounting cells on the bench at room temperature (∼5 min of preparation time) versus preparing cells inside the temperature-controlled cage installed on the microscope. For experiments at permissive or semi-permissive temperatures (Fig. S1), cells were maintained at a steady temperature (25 or 28°C) for ∼18 hr and imaged at the indicated temperature in the incubator. For inhibition of *cdc2-as* and *sid2-as* alleles (Fig. 1, S1, 2, S2), cells were grown in liquid EMM3S at 30°C with shaking and treated with 10µM 1-NM-PP1 (100-fold dilution of a 4mM stock in DMSO) (#50-203-0494, #67-68-5, Fisher Scientific) for 3-6 hr. Cells were harvested and imaged as above.

### Preparation of cells for live cell microscopy

Cells were placed just before imaging into μSlide VI 0.4 channel slides (#80606, Ibidi – 6 channels slide, channel height 0.4mm, length 17mm, and width 3.8mm, tissue culture treated and sterilized). The μSlide was first pre-coated by incubation with 100μg/mL of lectin (#L1395, Sigma) for at least 15 min at room temperature and then removed from the chamber. For mounting cells, 1 mL of liquid yeast culture was centrifuged for 2 min in an microcentrifuge tube at 400 x G at room temperature. Most supernatant was removed, and the cell pellet was gently resuspended in the remaining ∼100 µL media. 50 µL of this concentrated cell mixture was added to the pre-coated chamber and allowed to adhere for 2 min and washed three times with pre-warmed media to remove non-adhered cells.

### Microscopy

For imaging of cytGEMs (Fig. 1 and 2.3), live cells were imaged with a TIRF Diskovery system (Andor) with a Ti-Eclipse 2 inverted microscope stand (Nikon Instruments), a 488nm laser illumination, a 60X TIRF oil objective (NA: 1:49, oil DIC N2) (#MRD01691, Nikon), and a sCMOS camera (Zyla, Andor). These components were controlled with μManager v. 1.41 (Edelstein *et al*., 2010; Edelstein *et al*., 2014). Temperature was maintained by a black panel cage incubation system (#748-3040, OkoLab). Cells were mounted in μSlide VI 0.4 channel slides (#80606, Ibidi – 6 channels slide, channel height 0.4mm, length 17mm, and width 3.8mm, tissue culture treated and sterilized).

For imaging of nuclei and fluorescence intensity quantification (Fig. 2 and 2.5), cells were imaged on a Ti-Eclipse inverted microscope (Nikon Instruments) with a spinning-disk confocal system (Yokogawa CSU-10) that includes 488nm and 541nm laser illumination (with Borealis) and emission filters 525±25nm and 600±25nm respectively, 40X (NA: 0.6) and 60X (NA: 1.4) objectives, and an EM-CCD camera (Hamamatsu, C9100-13). These components were controlled with μManager v. 1.41 (Edelstein *et al*., 2010, 2014). Temperature was maintained by a black panel cage incubation system (#748–3040, OkoLab).

### Imaging and analysis of cytGEMs

Cells expressing cytGEMs nanoparticles were imaged in fields of 1K x 1.2K pixels or smaller using highly inclined laser beam illumination at 100Hz for 5 s. Cells generally exhibited 10-20 of cytGEMs nanoparticles/cell. CytGEMs were tracked with the ImageJ (Schindelin *et al*., 2012) particle Tracker 2D-3D tracking algorithm from MosaicSuite (Sbalzarini and Koumoutsakos, 2005) with the following parameters: run(“Particle Tracker 2D/3D”, “radius = 3 cutoff = 0 per/abs = 0.03 link = 1 displacement = 6 dynamics = Brownian”). In Figures 1-2, cytGEMs were analyzed collectively in multiple cells in the whole field of view. For analyses of individual cells (Fig. 1 D), cells were individually cropped from field images, and cytGEMs were tracked with the same MosaiSuite parameters with the exception of per/abs = 0.03. The analyses of the cytGEMs tracks were as described in Delarue *et al*., 2018, with methods to compute mean square displacement (MSD) using MATLAB (MATLAB_R2018, MathWorks). The effective diffusion coefficient D_eff_ was obtained by fitting the first 10 time points of the MSD curve (MSD_truncated_) to the canonical 2D diffusion law for Brownian motion: MSD_truncated_(τ) = 4 ⋅ Deff ⋅ τ.

### Measurement of cell length and nuclei count

As a proxy for cell size, cell length along the long axis of the rod-shaped cells was measured manually using ImageJ Line Selection tool on brightfield images of cells. “Straight Line” or “Segmented Line” was used depending on cell morphology. For determination of the number of nuclei, strains with the nuclear envelope marker Ish1 tagged with a fluorescent protein were grown in EMM3S (minus leucine) media, and number of nuclei were counted manually. Septated cells were excluded from analysis.

### Ribosomal concentration quantification

Ribosomal concentration was measured in individual fission yeast cells using the fluorescence intensity of ribosomal protein Rps2-GFP, as described (Knapp *et al*., 2019; Lemière *et al*., 2022). Cells expressing Rps2-GFP were grown in rich YES liquid media at 25°C overnight and shifted to 36°C for 6 hr before imaging. Cells were mounted on a 2% agarose (#16500500, Invitrogen) in YES 225 (#2011, Sunrise Science Production) pad and imaged with 488 nm laser illumination via spinning disk confocal microscopy. The Rps2-GFP signal was acquired in 500 nm z-step stacks, and a sum of stack of the middle 3 slices was used for intensity quantification. For each selected cell, the Rps2-GFP signal intensities were measured along the long cell axis (averaged over 4 µm in width) and normalized by cell length. The signal was corrected for background intensity and uneven illumination of the field. Rps2-GFP signals were defined as the average of the mean signal between 0.2-0.3 and the mean signal between 0.7–0.8 (peak signals in the cytoplasm, avoiding the nucleus) along the normalized cell length. Finally, all Rps2-GFP signals were normalized to the mean of the cell length (L) category 9 ≤ L < 18 µm.

### FITC staining to quantify cellular protein concentration

Total protein was measured in individual fission yeast cells using FITC staining, similar to as described (Knapp *et al*., 2019; Odermatt *et al*., 2021; Lemière *et al*., 2022). Cells were grown in YES liquid media at 25°C overnight and shifted to 36°C for 6 hr until fixation. 1 mL of exponential-phase (OD_600_ = 0.2-0.6) cell culture was fixed with 4% final concentration of formaldehyde (methanol-free 37% solution, #28906, Thermo Scientific, Waltham) and incubated at 4°C overnight. Fixed cells were washed 3 times with phosphate buffered saline (PBS) (#14190, Thermo Scientific) and resuspended in 100 µL of PBS. 100 µL of fixed cells were treated with 0.1 mg/mL RNAse A (#EN0531, Thermo Scientific) and incubated in a rotator for 2 hr at 37°C. Next, cells were washed and re-suspended in PBS and stained with 50 ng/mL FITC (#F7250, Sigma) for 30 min, washed three times with PBS, and resuspended in PBS. Cells were mounted on a 2% agarose (#16500500, Invitrogen) in Dulbecco’s Phosphate Buffer Saline (Thermo Scientific, 14190144) pad and imaged with 488 nm laser illumination via spinning disk confocal microscopy. FITC signal was acquired and analyzed using similar methods as the Rps2-GFP experiments described above.

### LC-MS/MS sample preparation

Proteomic experiments were performed using stable isotope labeling by amino acids in cell culture (SILAC) (Ong *et al*., 2002). SILAC-compatible fission yeast strains containing *car2Δ* were grown in SILAC adjusted media (Edinburgh Minimal Media (#4110712, MP Biomedicals) + 6 mM ammonium chloride + 0.04 mg/ml arginine and 0.03 mg/ml lysine) using either light or ‘‘heavy’’ versions of Lysine and Arginine (Swaffer *et al*., 2016). The ‘‘light’’ (Agr0 Lys0) version of the media contained L-Arginine and L-Lysine built with normal 12C and 14N isotopes; the ‘‘heavy” (Arg6 Lys4) version had L-Arginine containing six 13C atoms and L-Lysine containing four deuterium atoms. For SILAC experiments, cells were grown for at least 8 generations at the indicated temperatures 25-36°C with shaking before collection, diluted in the morning and evening so they are always below OD_600_ = 0.3. The mean cell volume for proteomics samples was determined by Z2 Coulter Counter (Beckman Coulter), and the mean cell volumes of these samples matched those of the corresponding samples used in the cytGEMs experiments.

10 mL of fission yeast cultures were pelleted by centrifugation at 3,000 x G for 2 min at 4°C. The supernatant was removed, and cell pellets were snap frozen in liquid nitrogen and stored at -80°C. Frozen pellets were resuspended in 300 µL of yeast lysis buffer (50 mM Tris, 150 mM NaCl, 5 mM EDTA, 0.2% Tergitol, pH 7.5; + a cOmplete ULTRA Tablet) with 700 µL of glass beads. Lysis was performed at 4°C in a MPBio Fastprep24 (4 cycles with the following settings: 6.0 m/s, 40 s). Cell lysates were cleared by centrifugation at 12,000 x G for 5 min at 4°C. Protein concentration was quantified using a Pierce BCA Protein Assay Kit (Prod# 23255). Equal amounts of protein from each SILAC-labeled lysate were mixed. The mixed lysates were then denatured/reduced in 1% SDS and 10 mM DTT (15 min at 65°C), alkylated with 5mM iodoacetamide (15 min at room temperature), and then precipitated with three volumes of a solution containing 50% acetone and 50% ethanol (on ice for 10 min). Proteins were re-solubilized in 2 M urea, 50 mM Tris-HCl, pH 8.0, and 150 mM NaCl, and then digested with TPCK-treated trypsin (50:1) overnight at 37°C. Trifluoroacetic acid and formic acid were added to the digested peptides for a final concentration of 0.2% (pH ∼3). Peptides were desalted with a Sep-Pak 50 mg C18 column (Waters). The C18 column was conditioned with 500 µL of 80% acetonitrile and 0.1% acetic acid and then washed with 1mL of 0.1% trifluoroacetic acid. After samples were loaded, the column was washed with 2 mL of 0.1% acetic acid followed by elution with 400 µL of 80% acetonitrile and 0.1% acetic acid. The elution was dried in a Concentrator at 45°C.

### LC-MS/MS data acquisition

Desalted SILAC-labeled peptides were analyzed on a Fusion Lumos mass spectrometer (Thermo Fisher Scientific, San Jose, CA) equipped with a Thermo EASY-nLC 1200 LC system (Thermo Fisher Scientific, San Jose, CA). Peptides were separated by capillary reverse phase chromatography on a 25 cm column (75 µm inner diameter, packed with 1.6 µm C18 resin, AUR2-25075C18A, Ionopticks, Victoria Australia). Peptides were introduced into the Fusion Lumos mass spectrometer using a 125 minute-stepped linear gradient at a flow rate of 300 nL/minute. The steps of the gradient are as follows: 3–27% buffer B (0.1% (v/v) formic acid in 80% acetonitrile) for 105 min, 27-40% buffer B for 15 min, 40-95% buffer B for 5 min, and finally maintained at 90% buffer B for 5 min. Column temperature was maintained at 50°C throughout the procedure. Xcalibur software (Thermo Fisher Scientific) was used for the data acquisition and the instrument was operated in data-dependent mode. Advanced peak detection was enabled. Survey scans were acquired in the Orbitrap mass analyzer (Profile mode) over the range of 375 to 1500 m/z with a mass resolution of 240,000 (at 200 m/z). For MS1, the Normalized AGC Target (%) was set at 250 and max injection time was set to “Auto”. Selected ions were fragmented by Higher-energy Collisional Dissociation (HCD) with normalized collision energies set to 31, and the fragmentation mass spectra were acquired in the Ion trap mass analyzer with the scan rate set to “Turbo”. The isolation window was set to 0.7 m/z window. For MS2, the Normalized AGC Target (%) was set to “Standard” and max injection time was set to “Auto”. Repeated sequencing of peptides was kept to a minimum by dynamic exclusion of the sequenced peptides for 30 s. Maximum duty cycle length was set to 1 s.

### Spectral searches

All raw files were searched using the Andromeda engine (Cox et al., 2011) embedded in MaxQuant (v1.6.7.0) (Cox and Mann, 2008). In brief, 2-label SILAC search was conducted using MaxQuant’s default Arg6/10 and Lys4/8. Variable modifications included oxidation (M) and protein N-terminal acetylation, and carbamidomthyl (C) was a fixed modification. The number of modifications per peptide was capped at 5. Digestion was set to tryptic (proline-blocked). Peptides were ‘‘Re-quantified’’, and maxquant’s match-between-runs feature was not enabled. Database search was conducted using the UniProt proteome - UP000002485. Minimum peptide length was 7 amino acids. FDR was determined using a reverse decoy proteome (Elias and Gygi, 2007).

### Peptide quantitation

Our SILAC analysis utilized MaxQuant’s ‘‘proteinGroups.txt’’ output file. Contaminant and decoy peptide identifications were discarded. When applicable, the “Leading Razor Protein” designation was used to assign non-unique peptides to individual proteins. Normalized SILAC ratios were used to determine changes in the relative concentrations of individual proteins.

### Protein annotations

Protein annotations in Figure 4 were sourced from UniProt columns named “Gene Ontology IDs” “Subcellular localization [CC]” or PomBase “Complex Annotations” unless otherwise noted (The Uniprot Consortium, 2022; Rutherford *et al*., 2024). Protein localization was strictly parsed so that each annotated protein belongs to only one of the designated groups. Proteins with 2 or more annotations were ignored (except for the “Cytoplasm/Nucleus” category which required a nuclear and cytoplasmic annotation and for categories, e.g. Histone, Chromosome, Nucleolus, which also contained a “Nucleus” annotation).

### 2D Annotation Enrichment analysis

Annotation enrichment analysis was performed as described previously (Cox and Mann, 2012). The protein annotation groups were deemed significantly enriched and plotted if the Benjamini-Hochberg FDR was smaller than 0.02. The position of each annotation group on the plot is determined by the enrichment score (S). The enrichment score is calculated from the rank ordered distribution of Protein Slope values:

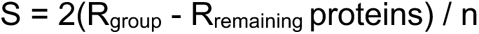

Where R_group_ and R_remaining_ proteins are the average ranks for the proteins within an annotation group and all remaining proteins in the experiment, respectively, and n is the total number of proteins. Highlighted annotation groups were manually curated.

### Gene ontology enrichment analysis

Relative protein concentration ratios were averaged between the two repetitions of proteomics experiments. Under- and overrepresented proteins were defined as having a minimum of a 10% change in their mean relative protein concentration ratio. GO process characterization of protein lists was performed using Protein Analysis Through Evolutionary Relationships (PANTHER) overrepresentation analysis version PANTHER 18.0 (Thomas *et al*., 2022).

## Acknowledgments

We thank the members of the Chang lab (present and past) for their generous support and discussion, and Joël Lemière for construction of ish1-tagged yeast strains. CT was supported by the National Science Foundation Graduate Research Fellowship Program (Award Number 1650113). FC was supported by National Institutes of Health grant GM141796. JS was supported by NIH R35 GM134858. MCL and the Mass Spectrometry were supported by a Collaborative Postdoctoral Fellowship with CZ Biohub San Francisco. MS was supported by a Wellcome Trust Career Development Fellowship. This work conducted as part of the first author’s dissertation work at UCSF (Tan, 2024).

## REFERENCES

1. Amodeo, AA, Jukam, D, Straight, AF, and Skotheim, JM (2015). Histone titration against the genome sets the DNA-to-cytoplasm threshold for the Xenopus midblastula transition. Proc Natl Acad Sci U S A 112, E1086–E1095.

2. Aoi, Y, Kawashima, SA, Simanis, V, Yamamoto, M, and Sato, M (2014). Optimization of the analogue-sensitive Cdc2/Cdk1 mutant by in vivo selection eliminates physiological limitations to its use in cell cycle analysis. Open Biol 4, 140063.

3. Balachandra, S, Sarkar, S, and Amodeo, AA (2022). The Nuclear-to-Cytoplasmic Ratio: Coupling DNA Content to Cell Size, Cell Cycle, and Biosynthetic Capacity. Annu Rev Genet 56, 165–185.

4. Basier, C, and Nurse, P (2023). The cell cycle and cell size influence the rates of global cellular translation and transcription in fission yeast. EMBO J 42, e113333.

5. Cadart, C, and Heald, R (2022). Scaling of biosynthesis and metabolism with cell size. Mol Biol Cell 33, pe5.

6. Chadwick, WL, Biswas, SK, Bianco, S, and Chan, YHM (2020). Non-random distribution of vacuoles in Schizosaccharomyces pombe. Phys Biol 17, 065004.

7. Chen, D, Toone, WM, Mata, J, Lyne, R, Burns, G, Kivinen, K, Brazma, A, Jones, N, and Bähler, J (2003). Global transcriptional responses of fission yeast to environmental stress. Mol Biol Cell 14, 214–229.

8. Claude, K-L, Bureik, D, Chatzitheodoridou, D, Adarska, P, Singh, A, and Schmoller, KM (2021). Transcription coordinates histone amounts and genome content. Nat Commun 12, 4202.

9. Cox, J, and Mann, M (2008). MaxQuant enables high peptide identification rates, individualized p.p.b.-range mass accuracies and proteome-wide protein quantification. Nat Biotechnol 26, 1367–1372.

10. Cox, J, and Mann, M (2012). 1D and 2D annotation enrichment: a statistical method integrating quantitative proteomics with complementary high-throughput data. BMC Bioinformatics 13 Suppl 16.

11. Creanor, J, and Mitchison, JM (1982). Patterns of protein synthesis during the cell cycle of the fission yeast Schizosaccharomyces pombe. J Cell Sci 58, 263–285.

12. Delarue, M, Brittingham, GP, Pfeffer, S, Surovtsev, I V., Pinglay, S, Kennedy, KJ, Schaffer, M, Gutierrez, JI, Sang, D, Poterewicz, G, et al. (2018). mTORC1 Controls Phase Separation and the Biophysical Properties of the Cytoplasm by Tuning Crowding. Cell 174, 338–349.e20.

13. Demidenko, ZN, and Blagosklonny, M V. (2008). Growth stimulation leads to cellular senescence when the cell cycle is blocked. Cell Cycle 7, 3355–3361.

14. Edelstein, A, Amodaj, N, Hoover, K, Vale, R, and Stuurman, N (2010). Computer control of microscopes using μManager. Curr Protoc Mol Biol. Chapter 14: Unit 14.20.

15. Edelstein, AD, Tsuchida, MA, Amodaj, N, Pinkard, H, Vale, RD, and Stuurman, N (2014). Advanced methods of microscope control using µManager software. J Biol Methods 1, e10.

16. Elias, JE, and Gygi, SP (2007). Target-decoy search strategy for increased confidence in large-scale protein identifications by mass spectrometry. Nat Methods 4, 207–214.

17. Elliott, SG (1983). Coordination of growth with cell division: Regulation of synthesis of RNA during the cell cycle of the fission yeast Schizosaccharomyces pombe. Molecular & General Genetics 192, 204–211.

18. Elliott, SG, Warner, JR, and McLaughlin, CS (1979). Synthesis of ribosomal proteins during the cell cycle of the yeast Saccharomyces cerevisiae. J Bacteriol 137, 1048–1050.

19. Facchetti, G, Knapp, B, Flor-Parra, I, Chang, F, and Howard, M (2019). Reprogramming Cdr2-Dependent Geometry-Based Cell Size Control in Fission Yeast. Current Biology 29, 350–358.e4.

20. Fantes, PA, and Nurse, P (1978). Control of the timing of cell division in fission yeast. Cell size mutants reveal a second control pathway. Exp Cell Res 115, 317–329.

21. Forsburg, SL (2003). Growth and Manipulation of S. pombe. In: Current Protocols in Molecular Biology, Chapter 13: Unit 13.16.

22. Garner, RM, Molines, AT, Theriot, JA, and Chang, F (2023). Vast heterogeneity in cytoplasmic diffusion rates revealed by nanorheology and Doppelgänger simulations. Biophys J 122, 767– 783.

23. Ginzberg, MB, Kafri, R, and Kirschner, M (2015). On being the right (cell) size. Science (1979) 348, 1245075.

24. Grallert, A, Connolly, Y, Smith, DL, Simanis, V, and Hagan, IM (2012). The S. pombe cytokinesis NDR kinase Sid2 activates Fin1 NIMA kinase to control mitotic commitment through Pom1/Wee1. Nat Cell Biol 14, 738–745.

25. Hachet, O, Berthelot-Grosjean, M, Kokkoris, K, Vincenzetti, V, Moosbrugger, J, and Martin, SG (2011). A Phosphorylation Cycle Shapes Gradients of the DYRK Family Kinase Pom1 at the Plasma Membrane. Cell 145, 1116–1128.

26. Hagan, IM, Riddle, PN, and Hyams, JS Intramitotic Controls in the Fission Yeast Schizosaccharomyces pombe: The Effect of Cell Size on Spindle Length and the Timing of Mitotic Events. J Cell Biol 110, 1617–21.

27. Harris, LK, Theriot, JA, and Harris, LK (2018). Surface Area to Volume Ratio: A Natural Variable for Bacterial Morphogenesis. Trends Microbiol 26, 815–832.

28. Hecht, VC, Sullivan, LB, Kimmerling, RJ, Kim, DH, Hosios, AM, Stockslager, MA, Stevens, MM, Kang, JH, Wirtz, D, Vander Heiden, MG, et al. (2016). Biophysical changes reduce energetic demand in growth factor–deprived lymphocytes. Journal of Cell Biology 212, 439– 447.

29. Heimlicher, MB, Bächler, M, Liu, M, Ibeneche-Nnewihe, C, Florin, EL, Hoenger, A, and Brunner, D (2019). Reversible solidification of fission yeast cytoplasm after prolonged nutrient starvation. J Cell Sci 132, 231688.

30. Joyner, RP, Tang, JH, Helenius, J, Dultz, E, Brune, C, Holt, LJ, Huet, S, Müller, DJ, and Weis, K (2016). A glucose-starvation response regulates the diffusion of macromolecules. Elife 5.

31. Keifenheim, D, Sun, X-M, Mayhew, MB, Marguerat, S, and Rhind, N (2017). Size-Dependent Expression of the Mitotic Activator Cdc25 Suggests a Mechanism of Size Control in Fission Yeast. Current Biology 27, 1491–1497.

32. Knapp, BD, Odermatt, P, Rojas, ER, Cheng, W, He, X, Huang, KC, and Chang, F (2019). Decoupling of Rates of Protein Synthesis from Cell Expansion Leads to Supergrowth. Cell Syst 9, 434–445.

33. Lanz, MC, Zatulovskiy, E, Swaffer, MP, Zhang, L, Ilerten, I, Zhang, S, You, DS, Marinov, G, McAlpine, P, Elias, JE, et al. (2022). Increasing cell size remodels the proteome and promotes senescence. Mol Cell 82, 3255–3269.e8.

34. Lanz, MC, Zhang, S, Swaffer, MP, Ziv, I, Hernández Götz, L, Kim, J, Mccarthy, F, Jarosz, DF, Elias, JE, Skotheim, JM, et al. (2024). Genome dilution by cell growth drives starvation-like proteome remodeling in mammalian and yeast cells. Nat Struct Mol Biol. 10.1038/s41594-024-01353-z

35. Lemière, J, Real-Calderon, P, Holt, LJ, Fai, TG, and Chang, F (2022). Control of nuclear size by osmotic forces in Schizosaccharomyces pombe. Elife 11, 1–41.

36. Lengefeld, J, Cheng, C-W, Maretich, P, Blair, M, Hagen, H, McReynolds, MR, Sullivan, E, Majors, K, Roberts, C, Ho Kang, J, et al. (2021). Cell size is a determinant of stem cell potential during aging. Sci Adv 7, 271.

37. Lloyd, AC (2013). The Regulation of Cell Size. Cell 154, 1194–1205.

38. Marguerat, S, and Bähler, J (2012). Coordinating genome expression with cell size. Trends in Genetics 28, 560–565.

39. Marshall, WF (2020). Scaling of Subcellular Structures. Annu Rev Cell Dev Biol 36, 219–236.

40. Martin, SG, and Berthelot-Grosjean, M (2009). Polar gradients of the DYRK-family kinase Pom1 couple cell length with the cell cycle. Nature 459, 852–856.

41. Maundrell, K (1993). Thiamine-repressible expression vectors pREP and pRIP for fission yeast. Gene 123, 127–130.

42. Mitchison, JM (1957). THE GROWTH OF SINGLE CELLS. Exp Cell Res 13, 244–262.

43. Mitchison, TJ (2019). Colloid osmotic parameterization and measurement of subcellular crowding. Mol Biol Cell 30, 173–180.

44. Molines, AT, Lemière, J, Gazzola, M, Steinmark, IE, Edrington, CH, Hsu, CT, Real-Calderon, P, Suhling, K, Goshima, G, Holt, LJ, et al. (2022). Physical properties of the cytoplasm modulate the rates of microtubule polymerization and depolymerization. Dev Cell 57, 466–479.e6.

45. Moseley, JB, Mayeux, A, Paoletti, A, and Nurse, P (2009). A spatial gradient coordinates cell size and mitotic entry in fission yeast. Nature 459, 857–860.

46. Mu, L, Kang, JH, Olcum, S, Payer, KR, Calistri, NL, Kimmerling, RJ, Manalis, SR, and Miettinen, TP (2020). Mass measurements during lymphocytic leukemia cell polyploidization decouple cell cycle-And cell size-dependent growth. Proc Natl Acad Sci U S A 117, 15659– 15665.

47. Munder, MC, Midtvedt, D, Franzmann, T, Nüske, E, Otto, O, Herbig, M, Ulbricht, E, Müller, P, Taubenberger, A, Maharana, S, et al. (2016). A pH-driven transition of the cytoplasm from a fluid-to a solid-like state promotes entry into dormancy. Elife 5, e09347.

48. Neumann, FR, and Nurse, P (2007). Nuclear size control in fission yeast. Journal of Cell Biology 179, 593–600.

49. Neurohr, GE, and Amon, A (2020). Relevance and Regulation of Cell Density. Trends Cell Biol 30, 213–225.

50. Neurohr, GE, Terry, RL, Lengefeld, J, Bonney, M, Brittingham, GP, Moretto, F, Miettinen, TP, Vaites, LP, Soares, LM, Paulo, JA, et al. (2019). Excessive Cell Growth Causes Cytoplasm Dilution And Contributes to Senescence. Cell 176, 1083–1097.e18.

51. Nurse, P (1975). Genetic control of cell size at cell division in yeast. Nature 256, 547–551.

52. Nurse, P (1985). Cell cycle control genes in yeast. Trends in Genetics 1, 51–55.

53. Nurse, P, Thuriaux, P, and Nasmyth, K (1976). Genetic control of the cell division cycle in the fission yeast Schizosaccharomyces pombe. Molecular & General Genetics 146, 167–178.

54. Odermatt, PD, Miettinen, TP, Lemière, J, Kang, JH, Bostan, E, Manalis, SR, Huang, KC, and Chang, F (2021). Variations of intracellular density during the cell cycle arise from tip-growth regulation in fission yeast. Elife 10, 1–23.

55. Ong, S-E, Blagoev, B, Kratchmarova, I, Kristensen, DB, Steen, H, Pandey, A, and Mann, M (2002). Stable Isotope Labeling by Amino Acids in Cell Culture, SILAC, as a Simple and Accurate Approach to Expression Proteomics*. Molecular and Cellular Proteomics 1, 376–386.

56. Padovan-Merhar, O, Nair, GP, Biaesch, AG, Mayer, A, Scarfone, S, Foley, SW, Wu, AR, Churchman, LS, Singh, A, and Raj, A (2015). Single Mammalian Cells Compensate for Differences in Cellular Volume and DNA Copy Number through Independent Global Transcriptional Mechanisms. Mol Cell 58, 339–352.

57. Pickering, M, Hollis, LN, D’Souza, E, and Rhind, N (2019). Fission yeast cells grow approximately exponentially. Cell Cycle 18, 869–879.

58. Rivas, G, and Minton, AP (2016). Macromolecular Crowding In Vitro, In Vivo, and In Between. Trends Biochem Sci 41, 970–981.

59. Rowley, R, Hudson, J, and Young, PG (1992). The wee1 protein kinase is required for radiation-induced mitotic delay. Nature 356, 353–355.

60. Rutherford, KM, Lera-Ramírez, M, and Wood, V (2024). PomBase: a Global Core Biodata Resource-growth, collaboration, and sustainability. Genetics 227, 1–14.

61. Sakai, K, Kondo, Y, Goto, Y, and Aoki, K (2024). Cytoplasmic fluidization contributes to breaking spore dormancy in fission yeast. Proc Natl Acad Sci U S A 121, e2405553121.

62. Sbalzarini, IF, and Koumoutsakos, P (2005). Feature point tracking and trajectory analysis for video imaging in cell biology. J Struct Biol 151, 182–195.

63. Schindelin, J, Arganda-Carreras, I, Frise, E, Kaynig, V, Longair, M, Pietzsch, T, Preibisch, S, Rueden, C, Saalfeld, S, Schmid, B, et al. (2012). Fiji-an Open Source platform for biological image analysis. Nat Methods 9, 676–682.

64. Schmoller, KM, Turner, JJ, Kõivomägi, M, and Skotheim, JM (2015). Dilution of the cell cycle inhibitor Whi5 controls budding-yeast cell size. Nature 526, 268–272.

65. Shi, H, Hu, Y, Odermatt, PD, Gonzalez, CG, Zhang, L, Elias, JE, Chang, F, and Huang, KC (2021). Precise regulation of the relative rates of surface area and volume synthesis in bacterial cells growing in dynamic environments. Nat Commun 12, 1–12.

66. Sun, XM, Bowman, A, Priestman, M, Bertaux, F, Martinez-Segura, A, Tang, W, Whilding, C, Dormann, D, Shahrezaei, V, and Marguerat, S (2020). Size-Dependent Increase in RNA Polymerase II Initiation Rates Mediates Gene Expression Scaling with Cell Size. Current Biology 30, 1217–1230.e7.

67. Swaffer, MP, Jones, AW, Flynn, HR, Snijders, AP, and Nurse, P (2016). CDK Substrate Phosphorylation and Ordering the Cell Cycle. Cell 167, 1750–1761.e16.

68. Swaffer, MP, Marinov, GK, Zheng, H, Fuentes Valenzuela, L, Tsui, CY, Jones, AW, Greenwood, J, Kundaje, A, Greenleaf, WJ, Reyes-Lamothe, R, et al. (2023). RNA polymerase II dynamics and mRNA stability feedback scale mRNA amounts with cell size. Cell 186, 5254–5268.e26.

69. Tai, YT, Fukuda, T, Morozumi, Y, Hirai, H, Oda, AH, Kamada, Y, Akikusa, Y, Kanki, T, Ohta, K, and Shiozaki, K (2023). Fission Yeast TORC1 Promotes Cell Proliferation through Sfp1, a Transcription Factor Involved in Ribosome Biogenesis. Mol Cell Biol 43, 675–692.

70. Tan, C (2024). Intracellular diffusion in the cytoplasm increases with cell size in fission yeast.

71. The Uniprot Consortium (2022). UniProt: the Universal Protein Knowledgebase in 2023. Nucleic Acids Res 51, 523–531.

72. Thomas, PD, Ebert, D, Muruganujan, A, Mushayahama, T, Albou, LP, and Mi, H (2022). PANTHER: Making genome-scale phylogenetics accessible to all. Protein Science 31, 8–22.

73. Urgen Cox, J, Neuhauser, N, Michalski, A, Scheltema, RA, Olsen, J V, and Mann, M (2011). Andromeda: A Peptide Search Engine Integrated into the MaxQuant Environment. J Proteome Res 10, 1794–1805.

74. Wood, E, and Nurse, P (2015). Sizing up to Divide: Mitotic Cell-Size Control in Fission Yeast. Annu Rev Cell Dev Biol 31, 11–29.

75. Xie, S, Swaffer, M, and Skotheim, JM (2022). Eukaryotic Cell Size Control and Its Relation to Biosynthesis and Senescence. Annu Rev Cell Dev Biol 38, 291–319.

76. Zatulovskiy, E, and Skotheim, JM (2020). On the Molecular Mechanisms Regulating Animal Cell Size Homeostasis. Trends in Genetics 36, 360–372.

77. Zhou, H-X, Rivas, G, and Minton, AP (2008). Macromolecular Crowding and Confinement: Biochemical, Biophysical, and Potential Physiological Consequences. Annu Rev Biophys 37, 375–397.

78. Zhurinsky, J, Leonhard, K, Watt, S, Marguerat, S, Bähler, J, and Nurse, P (2010). A coordinated global control over cellular transcription. Current Biology 20, 2010–2015.

